# Brain stimulation competes with ongoing oscillations for control of spike timing in the primate brain

**DOI:** 10.1101/2021.10.01.462622

**Authors:** Matthew R. Krause, Pedro G. Vieira, Jean-Philippe Thivierge, Christopher C. Pack

**Author notes:** These authors contributed equally (Matthew R. Krause, Pedro G. Vieira).

## Abstract

Transcranial alternating current stimulation (tACS) is commonly used to enhance brain rhythms, in the hopes of improving behavioral performance. Unfortunately, these interventions often yield highly variable results. Here, we identify a key source of this variability by recording from single neurons in alert non-human primates. We find that, rather than enhancing rhythmic activity, tACS appears to compete with the brain’s endogenous oscillations for control of spike timing. Specifically, when the strength of stimulation is weak relative to endogenous oscillations, tACS actually decreases the rhythmicity of spiking. However, when stimulation is comparatively stronger, tACS imposes its own rhythm on spiking activity. Thus the effect of tACS depends categorically on the strength of neural entrainment to endogenous oscillations, which varies greatly across behavioral states and brain regions. Without carefully considering these factors, attempts to impose external rhythms on specific brain regions may often yield precisely the opposite of the intended effect.

## Introduction

Transcranial electrical stimulation (tES) is a family of techniques that seek to modulate brain activity by applying electrical current non-invasively, through the scalp. This current flows through the head, producing electric fields that interact with the brain’s own electrical activity. By using tES with alternating current, researchers often attempt to target brain functions that rely on specific oscillation frequencies (1); this approach is called transcranial alternating current stimulation (tACS) (e.g., 2, 3). Although there have been some concerns about its effectiveness, there is now strong evidence that tACS can influence oscillatory neural activity in vitro (4, 5), in small animal models (6, 7), and even in the large, well-insulated primate brain (8–10).

Nevertheless, harnessing this mechanism to produce reliable changes in human behavior has proven to be surprisingly difficult. Studies often find that tACS produces inconsistent effects, between and within participants (e.g., 11, 12, 13), even with stimulation frequencies that are known to be linked to the specific behaviors under study. Although individual differences in neuroanatomy may explain some of these inconsistencies (12), variability in the participants’ ongoing brain activity also appears to shape the effects of tACS (14–18). For example, asking participants to close their eyes—which increases the amplitude of endogenous alpha oscillations—reduces the subsequent effects of tACS (17, 18). In contrast, increasing beta power, by imagining specific movements, seems to increase the effectiveness of tACS (16). The nature of the interactions between tACS and ongoing brain activity thus remains unclear. Since ongoing brain activity varies between (19, 20) and within (21, 22) individuals, understanding these interactions is critical for determining when, how, and for whom tACS will be effective.

Most behavioral studies are currently based on the assumption that tACS enhances ongoing oscillatory activity *(23)*, by causing neurons to fire in sync with the stimulation. Indeed, tACS is capable of creating subthreshold membrane potential fluctuations at the stimulation frequency *(24, 25)*, which might explain some reports that tACS and ongoing brain activity interact synergistically *(16).* At the same time, it has been argued that the relatively weak influence of tACS is likely to be overwhelmed by ongoing neural activity (*26–28*). If this were the case, tACS may be unable to alter the activity of neurons that are already entrained to an ongoing oscillation. These hypotheses can best be distinguished by measuring the influence of tACS on neurons with varying levels of entrainment to ongoing activity. However, prior neurophysiological experiments, including our own (*8–10*), have focused on conditions in which ongoing neuronal entrainment was weak. As a result, these experiments did not completely capture the conditions occurring during typical human tACS experiments.

Here, we characterize the interaction between ongoing oscillations and tACS, by recording single-neuron activity in the non-human primate brain. Our results confirm that tACS can entrain neural activity, but we find that this occurs only when spike entrainment to ongoing activity is weak. Surprisingly, when neurons are strongly locked to ongoing activity, applying tACS usually leads to a *decrease* in entrainment, which can only be reversed at higher stimulation amplitudes. Since the effects of tACS vary categorically with the strength of ongoing neural entrainment, our data indicate that it competes with brain oscillations for control over spiking activity. Moreover, we show that this competition is a straightforward mathematical consequence of interactions between oscillators.

These results have important ramifications for neuromodulation applications. On the one hand, the reduction of spike entrainment that we observe is precisely the opposite of the effect intended in most studies of human participants. On the other hand, we suggest that the same mechanism could be useful in attaining specific behavioral or clinical goals that require targeted desynchronization of neural activity. More generally, these findings also offer a possible mechanistic explanation for the extensive variability reported in the human tACS literature.

## Results

We examined the interplay of ongoing oscillations and tACS using recordings from non-human primates *(M. mulatta)*, a model system that captures many aspects of human anatomy, physiology, and tACS use. Animals were trained to perform a simple visual fixation task that minimized sensory and cognitive factors that influence oscillations. As animals performed this task, we recorded single-unit activity using standard neurophysiological techniques, as in previous work *(8, 9).* These data were then used to assess neuronal entrainment to ongoing local field potential (LFP) and tACS oscillations.

Our experiments targeted cortical area V4, where neurons often exhibit reliable entrainment to the (LFP), especially in the “theta” frequency band (*29*). We verified that this occurred in our experiments by computing phase-locking values (PLVs) that describe the consistency of spike timing (see *Materials and Methods).* These values range from zero (spiking occurs randomly across an oscillation’s cycle) to one (spiking occurs at only a single phase of the oscillation). We analyzed the entrainment of spikes to ongoing oscillations, by computing the PLV in 2 Hz frequency bins. In the absence of tACS, many V4 neurons were locked to the ~5 Hz component of the V4 LFP, with entrainment to higher frequency components being minimal (Figure S1). The strength of this entrainment varied across neurons, from 0 to 0.42, which provides an ideal way to test how tACS influences neural entrainment across different baseline entrainment conditions.

### tACS causes bidirectional changes in spike timing

Next, we recorded from the same V4 neurons during the application of tACS at 5 Hz. Our stimulation methods closely mimicked those commonly used in human studies and produced an electric field of similar strength (~ 1 V/m). The effects of tACS were assessed by comparing PLVs obtained during blocks of tACS against those computed from intervals of baseline or “sham” stimulation that were randomly interleaved as a control.

Data from four example neurons are shown in Figure 1A, which summarizes the phases at which spikes occurred during the baseline (blue) and tACS (orange) conditions. For these cells, the entrainment to the ongoing oscillation was weak, and the application of tACS led to a significant increase in phase-locking (p < 0.05, per-cell randomization tests; see *Materials and Methods).* These effects resemble those reported previously in other brain regions *(8–10)*, but they only occurred in 11% of the cells from which we recorded (17/157 neurons).

**Fig. 1.**
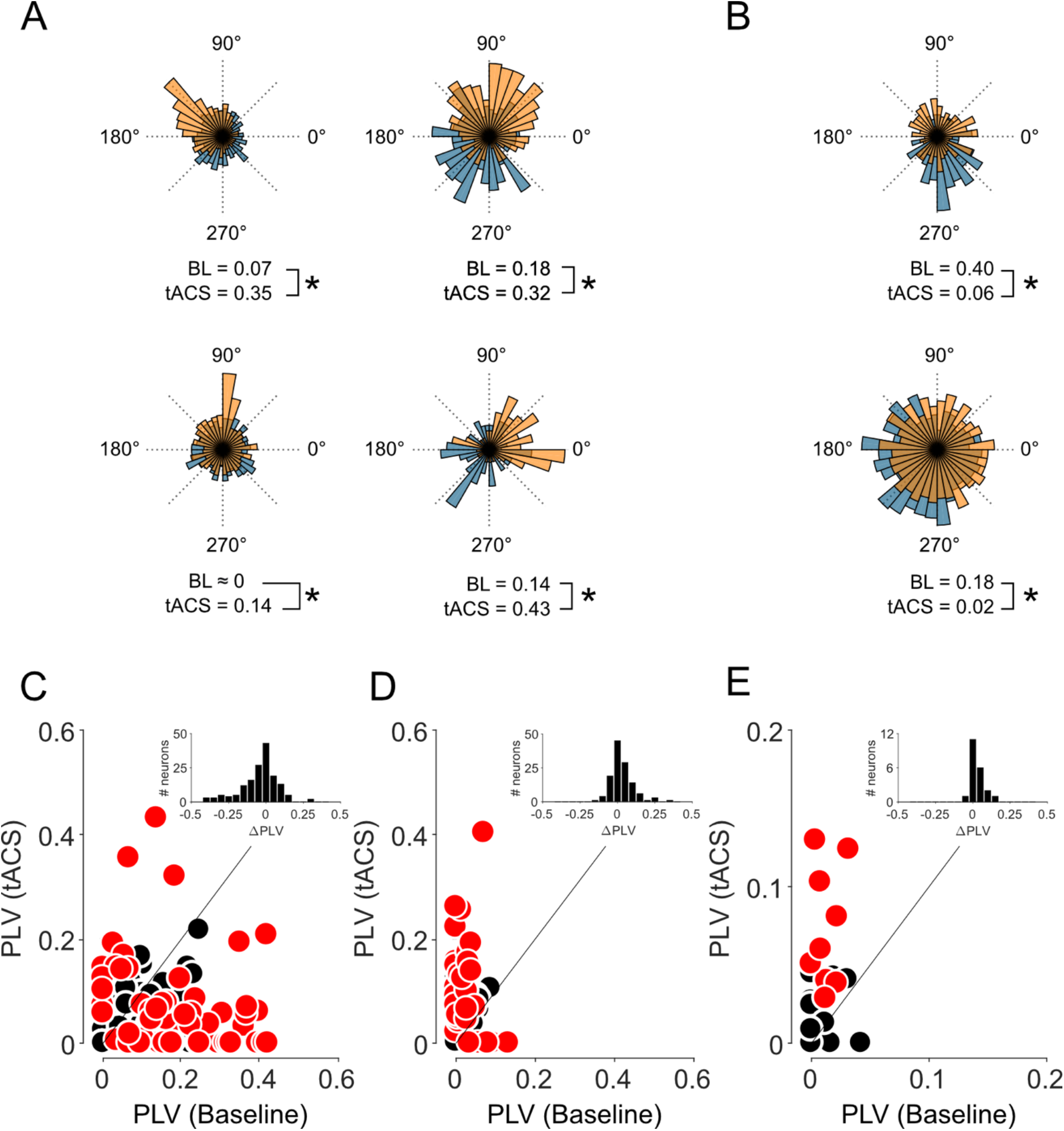
Applying tACS during physiological oscillations results in bidirectional changes in spike timing. **(A–B)** Spike-density histograms for six example V4 neurons showing the relative amounts of spiking across an oscillatory cycle. The cells in (A) showed increased entrainment, while those in (B) have decreased entrainment during 5 Hz tACS (orange) compared to the baseline LFP (blue, BL). PLV values for each condition are shown below; asterisks indicate p < 0.01. **(C–E)** Each point in the scatter plots represents a neuron’s PLV during baseline (horizontal position) and tACS (vertical). Data were collected (C) in the presence of a 5 Hz ongoing oscillation within V4 (N=157), (D) also in V4, but at 20 Hz, where the oscillation is weaker (N=123), and (E) again at 5 Hz but in the hippocampus (N=21), which also lacks strong 5 Hz oscillations under our conditions. Neurons showing individually significant changes in phase-locking are denoted in red (p < 0.05; per-cell randomization tests). Inset histograms show the changes in PLV across the population. Panel C adapted from Krause et al., (2019). See also Figure S1 for comparison of entrainment during baseline conditions.

For many other neurons, the application of tACS led to a *decrease* in the phase-locking of V4 neurons, as shown in the two example cells of Figure 1B. In these cases, the distribution of spike times was more uniform during tACS than during baseline. Statistically significant decreases in entrainment (p < 0.05; per-cell randomization tests) were found in 30% (47/157) of the neurons in our sample. Indeed, decreased entrainment was the predominant influence of 5 Hz tACS at the population level: the median PLV decreased from 0.066 (95% CI: [0.044 – 0.085]) to 0.031 [0.007 – 0.058], a statistically significant reduction in rhythmicity (p < 0.01; Z = 3.86; Wilcoxon sign-rank test). These changes were not accompanied by changes in firing rate: the median firing rate under baseline conditions was 4.6 Hz (95% CI: [3.7 – 5.8]) and 3.9 Hz (95% CI: [3.4—5.4]) during tACS, which was not significantly different (p > 0.1; Z = 1.63; Wilcoxon sign-rank). This suggests that these changes in entrainment were not due to signal loss or other artifacts (*9*).

Neurons which had higher levels of baseline entrainment tended to become less entrained during tACS, as can be seen in the example cells of Figure 1A-B. This pattern was evident in our complete dataset (Figure 1C), where baseline entrainment was significantly and negatively correlated with the subsequent changes during tACS (ρ = −0.38; p < 0.01). This correlation persisted even after the application of Oldham’s method (*30*) to guard against regression to the mean; a permutation-based analysis (*31*) yielded similar results.

### Decreased entrainment is not caused by stimulation frequency or brain region

The decreased entrainment we have observed may seem at odds with results from previous studies, which have consistently reported increased entrainment with tACS for spikes recorded from different brain regions and with different tACS frequencies *(6–10).* We therefore asked whether the lack of entrainment was due to our choice of brain region or stimulation frequency.

As a within-area control, we applied 20 Hz tACS instead. Neurons in V4 were weakly entrained to the 20 Hz LFP component under baseline conditions (median: 0.01; 95% CI: [0 – 0.018]), with only 1/123 neurons having a PLV above 0.1. Applying 20 Hz tACS to V4 caused 22 percent (27/123) of neurons to fire significantly more rhythmically (p < 0.05; per-cell randomization test; Figure 1B). The population PLV was significantly increased (p < 0.01, Z = −3.945; Wilcoxon sign-rank test), tripling from 0.01 to 0.029 [0 – 0.050]. These data demonstrate that V4 neurons can be entrained by tACS at a different frequency, suggesting that the 5 Hz results are not attributable to brain area.

As a within-frequency control, we also examined neural entrainment in the hippocampus, where baseline entrainment at 5 Hz was also weak. The median PLV was 0.01 and none of the 21 cells in our sample exhibited a PLV above 0.1. During 5 Hz tACS, entrainment increased in 9 of 21 hippocampal neurons (42%), and the median PLV significantly increased from 0.0083 [0 – 0.02] to 0.04 [0.01 – 0.05] (p < 0.01; Z=-3.06; Wilcoxon sign-rank test; Figure 1C). Together, these results show that decreased entrainment is neither a feature of V4 neurons nor of 5Hz tACS, both of which can be associated with increased entrainment under the right circumstances.

### tACS alters the strength and phase of entrainment

Some cells, such as the examples shown in the right column of Figure 1A, also showed a shift in the oscillatory *phase* at which spiking most often occurred. The specific timing of spikes within an oscillation (i.e., phase) may encode additional information (*32*), so shifts in preferred phases may represent another dimension along which tACS can alter neural activity. In principle, these changes could occur even when the overall levels of entrainment are not affected by tACS.

We therefore examined the subset of neurons whose PLVs were not significantly affected by tACS to see if their preferred phase of spiking shifted during stimulation. Since phase shifts require neurons to have a pre-existing phase preference, we first analyzed the 22/93 neurons which had individually-significant phase preferences (p<0.05; Rayleigh tests) under baseline conditions. Figure 2 plots the PLVs and preferred phases jointly for each neuron in each condition. The radial distance from the origin of corresponds to a neuron’s PLV, while the angular position denotes the phase preference. Vectors connecting the baseline values (blue dots) with those measured during tACS (grey dots) therefore completely describe the effects of stimulation on spike timing.

**Fig. 2.**
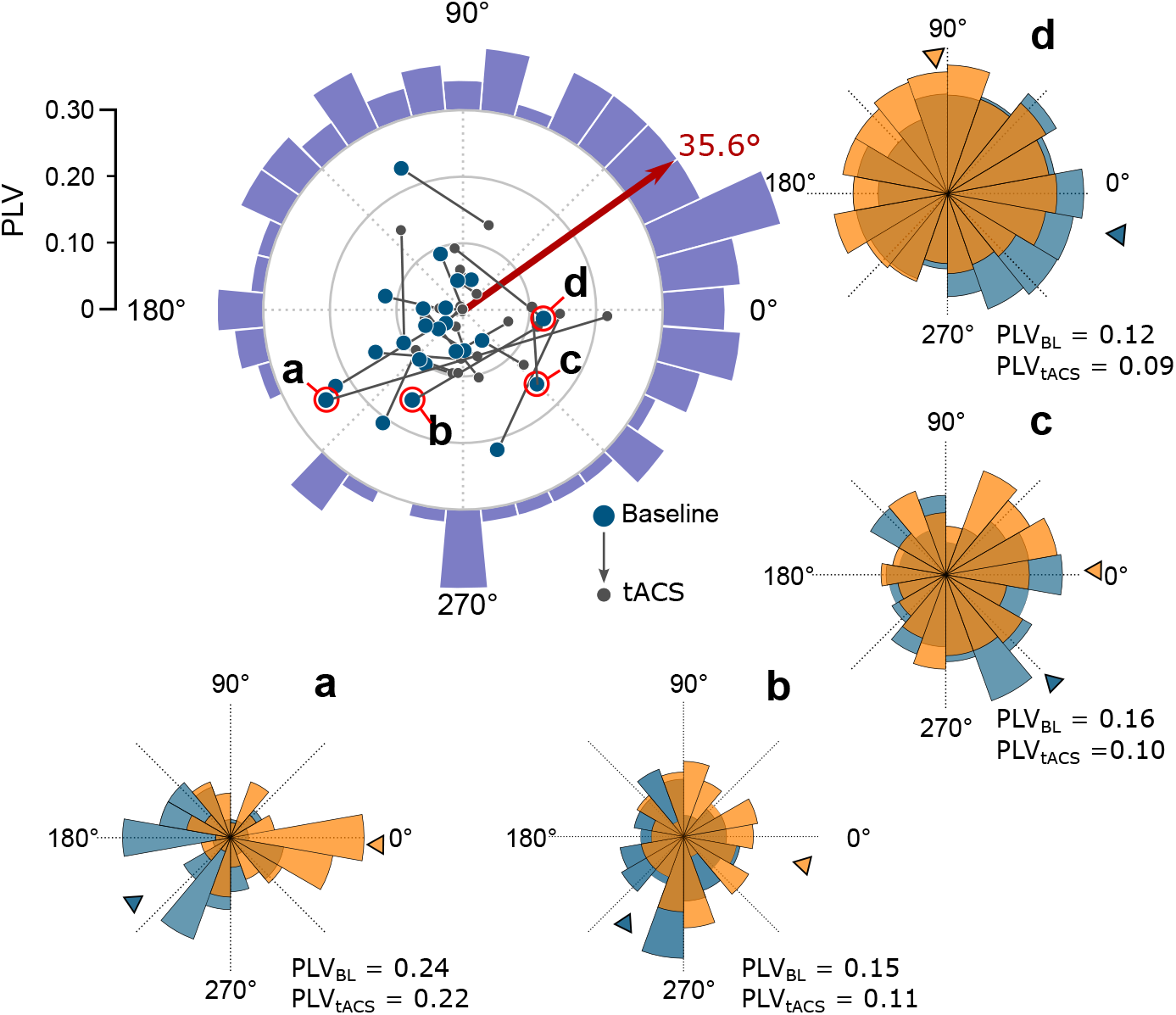
Ongoing oscillations determine the strength and phase of tACS entrainment. The polar plot summarizes the combined effects of tACS on entrainment strength (PLV, eccentric direction) and entrainment phase (polar angle). Each vector begins during baseline (blue dot) and extends to 1 mA tACS (grey dot) for the 22 neurons that had no significant change in PLV but a significant phase preference during baseline. Red arrow indicates their average direction of change. Spike-phase histograms are shown for the four example cells circled in red. The violet histogram shows similar changes in direction across the entire population of 93 neurons that whose PLVs were not significantly affected by tACS. It peaks at 38°, near the red arrow tip. See also Figure S2.

For these neurons, applying tACS consistently shifted the neurons’ spiking phase, so that they developed a statistically significant (p < 0.01; Hodges-Ajne omnibus test) preference for firing during the rising phase of the tACS waveform (35.6°, red arrow in Figure 2). This preference can be seen even in individual neurons, as demonstrated by the spike-phase histograms for the four example neurons in Figure 2. A broader analysis considering all 93 neurons that did not show altered entrainment during tACS revealed a similar shift towards 38° (p < 0.01; Hodges-Ajne omnibus test). The violet histogram wrapped around Figure 2 summarizes the change in phase for this larger population. These phase changes cannot be attributed to differences in referencing in the baseline and tACS conditions or to spike waveform distortions caused by tACS (see *Materials and Methods).*

We provide similar visualizations for the remaining neurons in Figure S2. Neurons that became entrained by tACS tend to start near the origin under baseline conditions and proceed outwards during stimulation (Figure S2A). Likewise, when tACS reduced a neurons’ entrainment to the ongoing oscillation, its spiking activity started in the periphery under baseline conditions and proceeded inwards (Figure S2B). We therefore suggest that the phase shifts shown in Figure 2 reflect a combination of these effects: tACS first decouples neurons from their entrainment to the ongoing oscillation, and then resynchronizes them to the stimulation waveform at a new phase.

### tACS first desynchronizes, then re-entrains neural activity

To test this possibility, we collected additional data from 47 V4 neurons, using both the original ±1 mA stimulation, as well as ±2 mA. We reasoned that if weak stimulation were partially reducing entrainment to a physiological oscillation, stronger electric fields could completely overcome it and lead to increased entrainment to the tACS waveform (*33*).

Figure 3 shows the results of this experiment. As in Figure 2, the location of each blue dot represents a neuron’s baseline entrainment, in terms of overall PLV (radial distance from the origin) and preferred phase (angle). Spike timing during ±1 mA is depicted by orange dots, and ±2 mA by red dots, forming a trajectory that shows the effects of increasing current on spike timing. Our hypothesis predicts that these trajectories should have a specific form: Starting from the neuron’s baseline level of entrainment, they first move towards the origin as the tACS and LFP vie for control of spike timing. Once tACS overwhelms the baseline entrainment, it imposes its own rhythm and the vector extends outwards from origin towards the rising phase of the tACS.

**Fig. 3.**
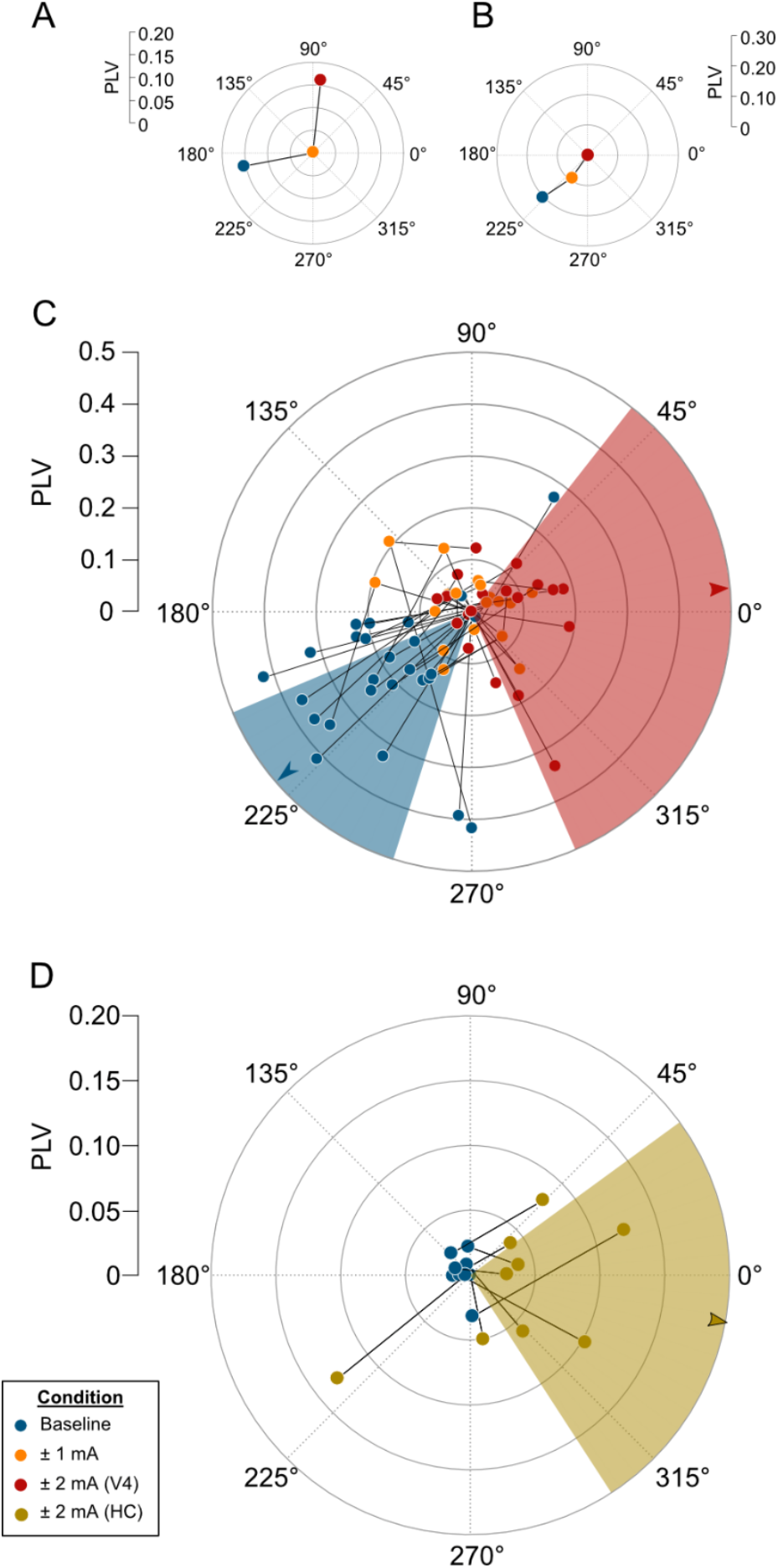
tACS reduces entrainment, then reinstates it at a different phase, as the stimulation amplitude increases. **(A-C)** Polar vectors indicating the phase and strength of V4 neurons’ entrainment (as in Figure 3. The blue dots indicate baseline conditions (blue), followed by ±1 mA tACS (orange) and ±2 mA tACS (red). Shading indicates the median and 95% confidence interval of the phase preference during baseline (blue) and ±2 mA tACS (red). Panels A and B show an example trajectory for one neuron exhibiting reinstated entrainment (A) and another undergoing progressive decreases in entrainment (B). Panel C shows population data from 27 neurons with significant changes in PLV at either amplitude. **(D)** Data from N = 9 hippocampal neurons during baseline (blue dot) and ±2 mA tACS (olive), plotted as in Panel A. Note that ±2 mA tACS produces an electrical field within the hippocampus that is three times weaker than the ±1 mA condition for V4.

Many neurons exhibited this pattern: applying ±1 mA tACS decreased or eliminated entrainment relative to baseline, but the stronger ±2 mA stimulation reinstated some entrainment, often at a different phase. The example neuron shown in Figure 3A had a PLV of 0.159 under baseline conditions, no detectable entrainment during ±1 mA tACS (PLV ≈ 0), and a PLV that again reached 0.158 during ± 2 mA stimulation. During this transition from 0 to ±2 mA stimulation, its preferred phase rotated from 191° to 83°. While a naïve analysis that considered only changes in PLV might conclude that this neuron was insensitive to tACS, these data demonstrate that the structure of its spike timing is, in fact, altered by the stimulation.

Other neurons instead progressively lost entrainment as the stimulation amplitude increased. This pattern appears in the example cell shown in Figure 3B, whose PLVs decreased from 0.21 to 0.092, to nearly 0 as the tACS amplitude increased, with the phase consistently remaining near 225° (223°, 235°, and 221°, respectively). These patterns are common in our data, as shown in Figure 3C, which depicts the trajectories through 0, ±1, and ±2 mA for the 28/47 neurons (60%) in our data set that showed significant changes from their baseline PLV at either tACS intensity. One possible interpretation of these data is that stimulation was less effective for these cells (e.g., due to their orientation relative to the electric field *(34))* and that stimulation above ±2 mA would be necessary to completely desynchronize and re-entrain them.

We present a similar diagram for our hippocampal data in Figure 3D. Although the electric field reaching this deep structure was far weaker (0.2—0.3 V/m; *9*), nearly half of these neurons (9/21; Figure 1C) showed increased entrainment, reminiscent of the ±2 mA condition in V4. Critically, these neurons’ baseline levels of entrainment were near zero, so it was not necessary to overcome any substantial baseline entrainment and stimulation can immediately entrain their spike timing. Interestingly, the preferred phase for these neurons was similar to that observed in V4 (Figure 3C). Overall, these results, across stimulation frequencies, stimulation amplitudes, and brain regions, show consistently that tACS influences spiking activity in a manner that depends strongly on the levels of pre-existing entrainment.

### Similar patterns of effects emerge from a simple oscillator model

Our experimental results show a specific pattern of reduced entrainment from baseline, followed by restored entrainment at higher stimulation amplitudes. Although there are many ways in which tACS can interact with ongoing oscillations (reviewed in *35*), most previous experimental work has not reported decreased entrainment. We therefore sought to determine whether decreased entrainment has a theoretical basis in the properties of coupled oscillators. Specifically, we explored the properties of a simple oscillator model that consists of two equations (*36*):

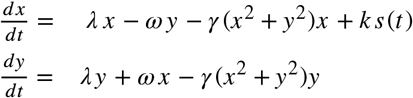

Here, *x* and *y* represent the dynamics of two coupled populations of neurons, whose interactions generate an oscillation. For simplicity, we assumed that higher amplitude oscillations were associated with stronger phase-locking, as found in previous experimental work *(7, 25).*

We simulated the effects of tACS by changing the properties of the external drive *s(t).* As in our experiments, *s(t)* was either zero (for baseline conditions) or a sine wave of given frequency and phase offset for the tACS condition. The coupling parameter *k* determines how strongly this external drive affects the neuronal population. In the results below, we express *k* as a percentage of the ongoing oscillation’s amplitude, which simplifies the model.

We first asked whether a consistent *phase* offset between an ongoing oscillation and tACS at the same frequency could account for our results. It did not. Stimulation applied at all phase offsets led to phase shifts in the ongoing oscillation, accompanied by an increase in its amplitude. For example, Figure 4A shows that applying even 180° out-of-phase stimulation of 5%, 30%, and 75% increased the oscillation’s amplitude by 5%, 26%, and 49% respectively, while shifting the phase by up to a quarter cycle relative to the no-stimulation condition.

**Fig. 4.**
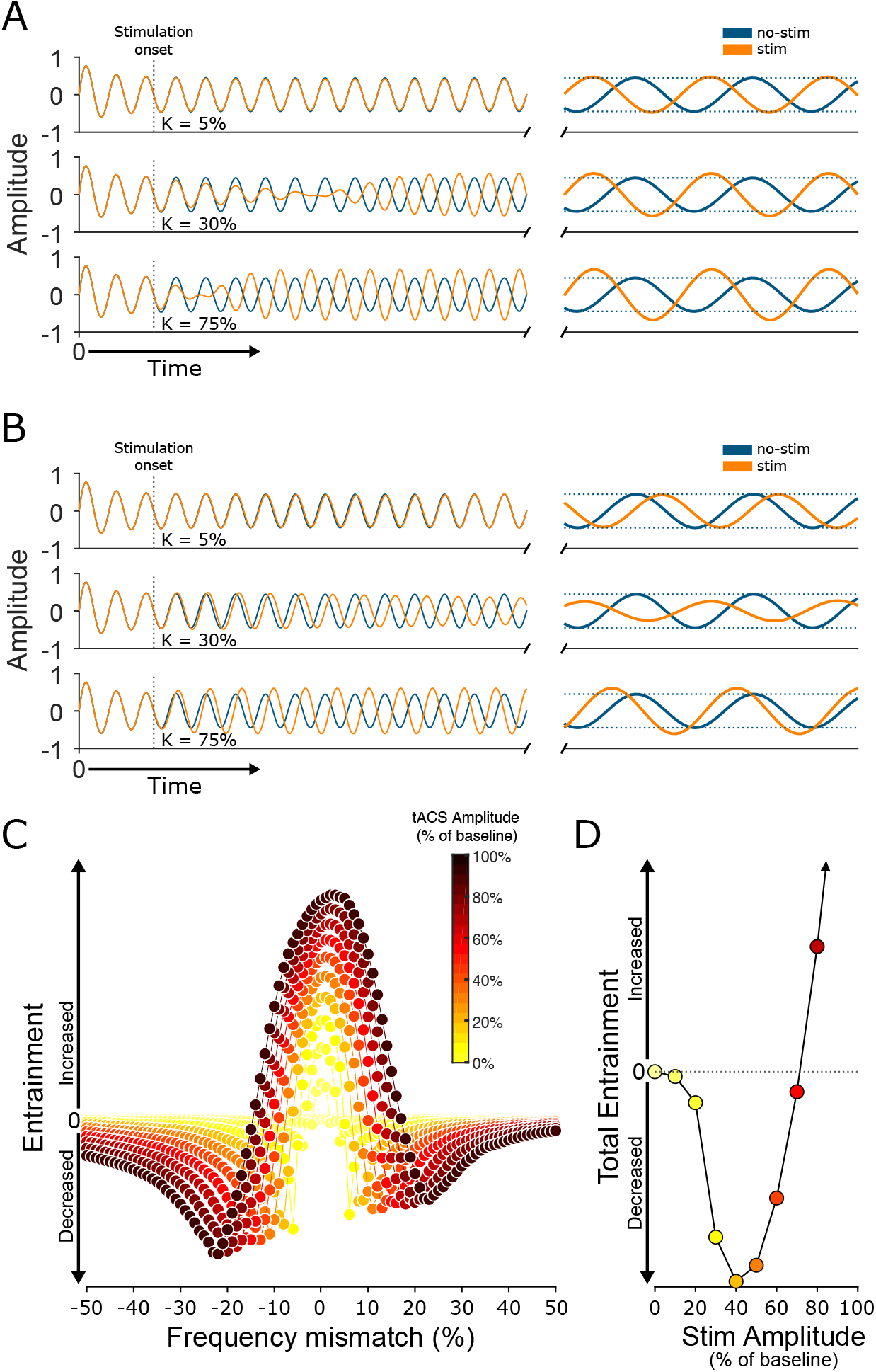
A simple oscillator model replicates these results. **(A)** Stimulating the Stuart-Landau model at the same frequency tends to increase entrainment and shift phase. The model’s output, *x(t)*, is shown during a no-stimulation condition (blue) and during the application of tACS (yellow) at low (top), medium (middle), and high (bottom) stimulation intensities. Dotted lines indicate oscillation’s amplitude in the absence of stimulation. The other component of the model, *y(t)*, is simply a phase-shifted version of *x(t)* and shows the same pattern of effects, as shown in Figure S3. **(B)** Stimulation applied at a slightly different frequency tends to reduce entrainment instead. The model output is plotted as in Panel A. **(C)** Changes in entrainment, as a function of frequency mismatch and stimulation intensity. **(D)** Changes in total entrainment, calculated by integrating the curves in Panel C. Colors correspond to those used in Panel C. See also Figures S4-S6.

We next asked whether slight differences between the *frequencies* of tACS and the ongoing oscillation could account for our results. This did seem to be the case: Figure 4B shows that applying stimulation at frequency that was only 6% below that of the ongoing oscillation led to an 29% decrease in amplitude at moderate stimulation intensities (k=30%; middle row of Figure 4B), followed by a 35% increase at higher intensities (k=75%; bottom row of Figure 4B). Stimulation that was instead slightly higher in frequency had similar effects, causing a 25% decrease, followed by a 44% increase (not shown) in the ongoing oscillation’s amplitude. Importantly, this effect on the strength of entrainment was independent of the relative phases of the tACS and ongoing oscillations. Figure 4C summarizes this effect for a range of relative frequencies, by averaging across relative phases, for consistency with the open-loop nature of our experiments. Stimulation that precisely matched the oscillation’s frequency led to increased oscillation strength but detuning the frequency by even a fraction of a cycle reduced the amplitude noticeably when the stimulation amplitude was relatively weak.

In real experiments, such a mismatch is likely unavoidable, given variability in spike timing (*37*) and peak frequencies within an oscillation (*38*), as well as the non-sinusoidal nature of neural oscillations (*39, 40*). Moreover, most analysis methods consider entrainment within a range of frequencies, as we have done here (using 2 Hz analysis bins; see Methods), so that tACS at a single frequency will elicit mixed effects on entrainment.

We therefore examined the net effect of stimulation on narrow-band oscillations, by integrating the curves of Figure 4C across frequencies for each stimulation intensity (Figure 4D). Initially, the total entrainment within these narrow frequency bands decreased, but when the stimulation amplitude exceeded about 66 percent of the ongoing oscillation’s amplitude, entrainment increased, as we have observed in our data (Figure 3). Measurements of ongoing membrane potentials (*41, 42*) indicate that they are often several times larger than polarization produced by tACS (*34*), so tACS, as applied in human experiments, may often lie near this transition.

### Baseline frequency preference determines the effects of tACS

A straightforward prediction of this model is that entrainment should be most strongly reduced for neurons with frequency preferences that are displaced from the stimulation frequency (i.e., in the flank of the entrainment vs. frequency-mismatch curve in Figure 4C). To test this prediction, we reanalyzed our V4 data to identify each neuron’s baseline frequency preference with greater precision (in ±0.25 Hz frequency bands from 2-8 Hz). We calculated the PLV within each bin of these narrowband signals and assigned the bin with the highest baseline PLV as the neuron’s preferred narrowband frequency (Figure 5A).

**Fig. 5.**
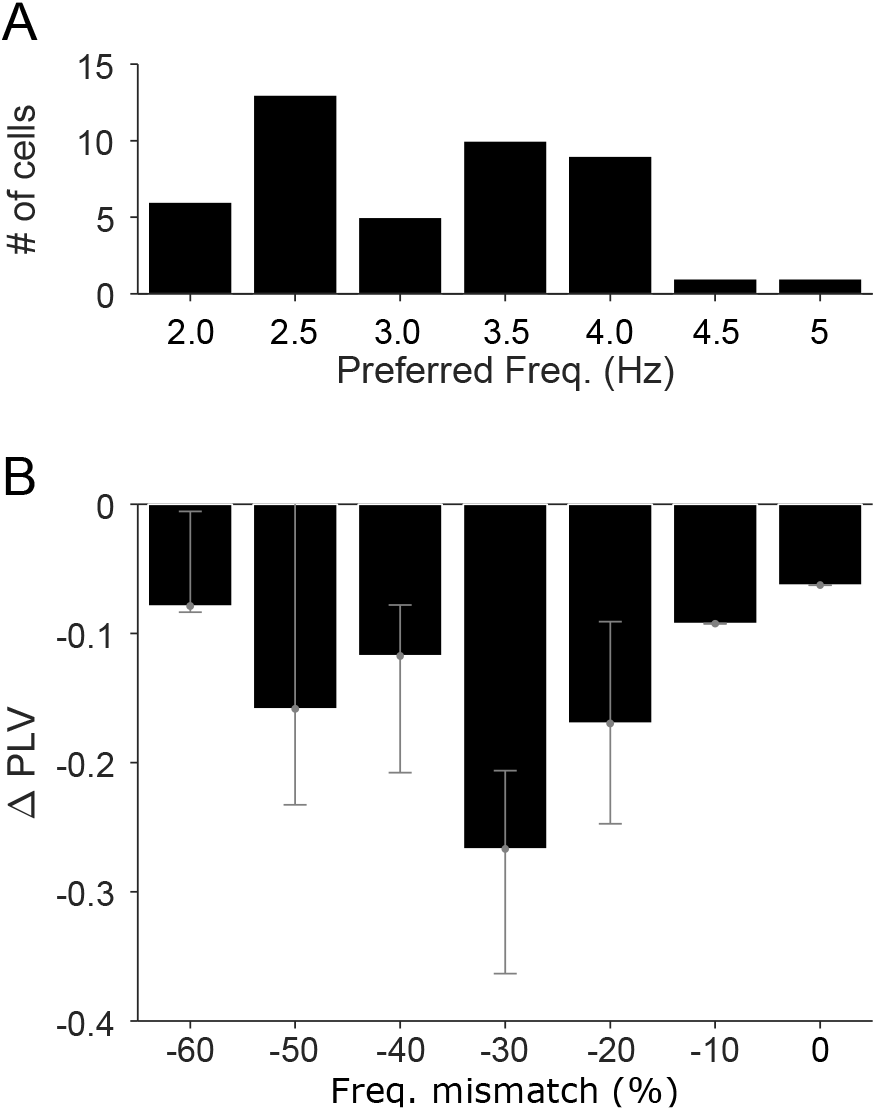
Neurons’ pre-existing frequency preference determines the effects of tACS. **(A)** Histogram of narrowband preferred frequency for the 47 V4 neurons with significantly decreased entrainment during 5 Hz tACS. **(B)** Changes in these neurons’ PLV as a function of the difference between their narrowband preferred frequency and the 5 Hz tACS frequency. Note the qualitative similarity to the flank of Figure 4C.

Figure 5B shows that, of the 47 neurons showing significantly decreased entrainment during 5 Hz tACS, most had baseline frequency preferences 1-2 Hz away from the stimulation frequency, reminiscent of the flank of Figure 4C. Only one neuron with decreased entrainment had a baseline frequency preference between 4.75–5.25 Hz. Owing to their low baseline entrainment, no pre-existing frequency preference could be identified for the 17 neurons that exhibited increased entrainment during tACS. These results demonstrate a strong but qualitative agreement with models of competing oscillators.

### Excitatory and inhibitory cells are similarly affected by tACS

The model makes another specific prediction that is of immediate biological relevance. Previous work has suggested that tACS may preferentially target certain types of neurons: due to their morphology, interneurons have been proposed to be less susceptible to tACS (*34, 43*). Others have suggested that, due to their positions with neural circuits, interneurons are actually more strongly affected by tACS (*44*). In the model, stimulation is applied to one subpopulation (the *x* term above), but both subpopulations are affected similarly by the stimulation (Figure S3), due to the tight coupling between them. This suggests that all neurons, regardless of cell type, will be affected similarly by tACS.

We therefore asked whether cell type could explain the mix of increased and decreased entrainment shown in Figure 1C. Putative cell types were identified via their spike widths: narrow-spiking neurons are often interneurons, while those with broader spikes are more likely to be pyramidal cells (See *Materials and Methods* and Figure 6A). As predicted from the model, broad-spiking neurons were no more likely than narrow-spiking neurons to be significantly modulated, in either direction, by tACS (p = 0.44; *X*^2^(1) = 0.59). When considering only the neurons affected by tACS (Figure 6B, colored dots), the two cell types did exhibit significantly different directions of modulation (p < 0.01; *X*^2^(1) = 12.1). Entrainment overwhelmingly decreased in broad-spiking neurons (88%; 36/41 cells), but the effects on narrow-spiking neurons were more mixed: 12 of 23 narrow-spiking cells became more entrained during 5 Hz tACS, while 11 of 23 showed significantly reduced entrainment. However, this appears to be driven by differences in baseline levels of entrainment: broad-spiking neurons tended to be strongly entrained at baseline, while narrow-spiking neurons fell into a weakly-entrained cluster with PLVs near 0 and a more strongly entrained one around 0.3 under baseline conditions. As in the full dataset, the effect of tACS on these narrow-spiking neurons was inversely correlated with baseline entrainment levels (Spearman’s ρ = −0.38; p=0.083). Data collected during 20 Hz tACS, where both cell types showed minimal baseline entrainment, and subsequently became strongly entrained by 20 Hz tACS (Figure 6C), further supports this hypothesis. Thus, the effects of tACS appear largely determined by a cell’s baseline level of entrainment, rather than cell type.

**Fig. 6.**
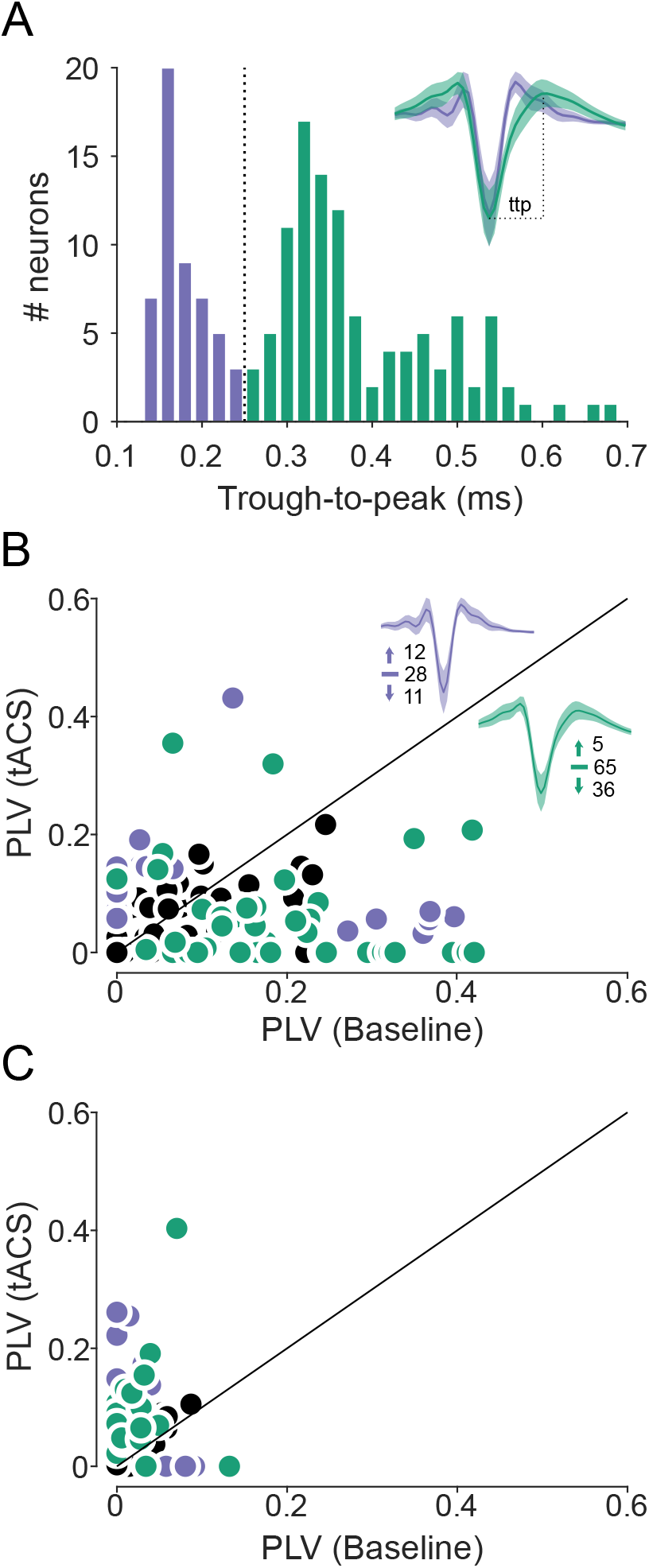
Distinct cell types are associated with tACS synchronization vs. desynchronization. **(A)** Neurons (N=157) were divided using a trough-to-peak width threshold of 250 μs (dashed line). Narrow-spiking putative interneurons are shown in purple; broad-spiking pyramidal cells in green. Average waveforms for each class are inset. **(B)** Changes in V4 neurons’ spike timing during 5 Hz tACS, color-coded by cell type; neurons without significant changes in PLV are shown in black. Numbers near the waveforms indicate the number of cells showing significantly increased, unchanged, and decreased PLVs. The style is otherwise identical to Figure 1A. **(C)** Changes in V4 neurons’ spike timing during 20 Hz tACS (N=123), plotted as in Panel B.

## Discussion

The results shown here demonstrate that tACS can cause individual neurons to fire either more or less rhythmically (Figure 1). The direction of these effects depends both on the relative strengths of entrainment to the ongoing physiological oscillation and the tACS-induced electric field (Figures 2 and 3). These data are consistent with a simple model (Figure 4) in which tACS competes with ongoing oscillations for control of spike timing, a hypothesis supported by several ancillary tests of the model (Figures 5 and 6). Producing predictable changes in neural activity, and ultimately, behavior, therefore requires detailed knowledge of the relevant oscillations within the targeted brain areas. Our limited understanding of these phenomena, even in the absence of stimulation, may be an underappreciated obstacle to the effective application of tACS. However, once these factors are understood and the stimulation tailored appropriately, the ability to increase and decrease the regularity of spike timing has numerous applications in the laboratory, clinic, and real world.

### Implications for human neuromodulation

The ability to increase and decrease neuronal entrainment opens up a number of new therapeutic possibilities. Excess synchrony has been implicated in a wide range of conditions, including movement disorders, epilepsy, and schizophrenia (*45*), and stimulation that reduces it may provide a means for controlling these symptoms. Deep Brain Stimulation (DBS) has become a standard, albeit invasive and expensive, method for treating severe cases of some of these diseases. While the mechanisms of action are not completely understood, it seems increasingly likely that DBS acts by regulating spike timing in ways that reinstate normal patterns of activity, rather than merely suppressing pathologically strong levels of activity (*46*). Our data suggests that tACS may be capable of producing similar effects non-invasively.

In power and aeronautical engineering, unwanted oscillations are sometimes controlled by applying open-loop stimulation at a frequency that is slightly “detuned” from that of the original oscillation. This approach, called “asynchronous quenching” or “amplitude death”, can be used in any system consisting of self-excited, forced oscillators, a category which includes the brain *(47).* Asynchronous quenching may therefore provide a simpler and more efficient way to regulate neural activity than closed-loop systems (*2*) that are often proposed for this task.

Traces of asynchronous quenching also appear in studies using rhythmic sensory stimulation. Listening to a 40 Hz tone suppresses, rather than enhances, gamma activity in auditory cortex (*48*), while visual flicker can also induce a combination of entrainment and event-related desynchronization (*49*).

At the same time, these data suggest that it may be much harder than expected to non-invasively increase neuronal entrainment to ongoing oscillations. To do so, stimulation must target brain regions and behavioral conditions in which ongoing entrainment is relatively weak, or else it must be precisely matched to both the frequency and phase of the on-going oscillation. The latter approach is challenging, because estimates of frequency and phase provide only transient approximations to the actual oscillations (*40*), which drift on sub-second timescales (*38, 39*). Thus, closed-loop stimulation may prove to be more useful for reinforcing existing oscillations.

However, most human tACS use is currently performed open-loop, using waveforms of a fixed frequency applied at an arbitrary time. Since those experiments typically seek to “increase”, “enhance”, (e.g., *50*) or “rescue” (*3*) an existing neural oscillation, our data raise the worrying possibility that many human tACS experiments have inadvertently produced the exact opposite of their intended effects. While the amplitude of endogenous oscillations varies considerably between areas and brain states, some normal oscillations shift membrane potentials by several millivolts (*42*). Since the direct effects of tACS impose polarizations of < 1 mV *(26, 34)*, this ratio clearly falls within the desynchronizing regime shown in Figure 4D.

### Implications for behavioral studies

In that light, we suggest some alternative interpretations of well-established tACS behavioral effects. Detection of faint auditory, visual, and somatosensory stimuli can often be improved by tACS (reviewed in *51*). These results are often ascribed to synchronization between the sensory input and the tACS waveform, as would be expected if tACS further entrained neurons that were already locked to sensory input. However, our data demonstrate that tACS often decreases neural entrainment under these conditions. Interestingly, decreased neural synchrony can sometimes lead to improved sensory performance, as in perceptual decision-making (*52, 53*) or selective attention (*54*). Some theories of attention propose that it improves visual performance by decreasing synchronization at lower frequencies, while simultaneously increasing it at higher ones (*55*). Since ongoing low-frequency oscillations tend to be strong, while entrainment to higher frequencies is often weak, a similar pattern of neural effects would seem to be an ideal way for tACS to improve performance on perceptual tasks. Similar mechanisms (*56, 57*) might account for improved sensory discrimination following transcranial magnetic stimulation, as suggested previously (*58*).

More generally, our results suggest that the effects of tACS likely depend on the behavioral task, insofar as many tasks modulate the strength of ongoing neural oscillations. The 8–12 Hz “alpha” frequency band is a very common target of tACS, but endogenous alpha oscillations vary in amplitude, depending on task demands (*21, 22*) and the overall alertness of the subject (*59*). Moreover, the dominant frequency within the alpha band varies across subjects, in a manner that is correlated with age and sex (*19*), clinical status (e.g., *20*), and other factors. As a result, the effects of fixed-frequency tACS (e.g., at 10 Hz) may vary across subjects as well as within a single subject as their performance of a task changes. Mechanisms like these may account for some of the extensive intra- and inter-subject variability observed in many tACS experiments (*12*).

Previous neurophysiological studies, including our own, have not found correlations between baseline activity and tACS’s effects on single neurons. We suggest that this is due to Simpson’s Paradox: the limited range of baseline PLVs in those studies masked the correlations reported here. However, our current findings are consistent with human tES experiments reporting similarly complex non-monotonic dose-response relationships on behavior (*60*) and motor-evoked potentials (*61*). Our modelling suggests that conditions within the human brain are often near the “elbow” of the curve shown in Figure 4D and that it may be advantageous to try multiple stimulation intensities. Although considerable effort has been devoted to maximizing the field strength produced by tACS montages, future studies should consider the possibility that, for some goals, more, in terms of current or electric field strength, may not always be better.

### Implications for biophysics

These data challenge previous work suggesting that tACS preferentially engages specific cell types. Based on their elongated, asymmetric morphology, pyramidal cells were predicted to be strongly susceptible to external electric fields while the smaller and bushier interneurons would be much less affected *(34, 43).* Although these biophysical predictions are likely valid for isolated neurons, they appear less relevant when the neurons are embedded in the complex, interacting networks of a living brain. Other work has suggested that network interactions cause tACS to produce especially strong effects in interneurons, which in turn drive the remaining neurons (*44*). Neither pattern is evident in our data (Figure 6), where cells’ ongoing patterns of activity, rather than morphology or identity, determine how they are affected by tACS. However, neurons’ entrainment to ongoing oscillations has been argued to reflect differences in their functional roles, such as their tendency to be affected by top-down input (*62*). The inverse correlation we observed may therefore affect these classes differently, and so the behavioral consequences of tACS may depend on more than the net change in entrainment across the entire neuronal population.

Previous work has also suggested that tACS affects only a small subset of neurons. For example, (*8*) reported that tACS alters PLVs in 9 to 27 percent of neurons, depending on the stimulation intensity. Likewise, *(7)* reported that less than 30 percent of neurons were entrained by fields of ~1 V/m, and our previous work also found tACS-induced PLV changes in only a third to a half of neurons (*9, 10*). However, these reports considered only individually significant net changes in PLV. The phase shifts demonstrated in Figure 2 suggest that tACS may affect many more neurons. Thus, experimental constraints and the choice of metrics has led the field to overestimate the sparseness of tACS’s effects.

Finally, several studies have reported that tACS increases EEG power (*63, 64*) which are often interpreted as evidence of successful neuronal entrainment by tACS. However, EEG signals likely arise from all transmembrane currents (*65*), including those which do not reach spike threshold. The two signals therefore measure slightly different aspects of neural information processing, and so these results are not necessarily incompatible with the ones we report here.

In summary, our results show that at the single neuron level, tACS competes with ongoing brain activity for control of spike timing. The net effect of this competition may be to increase or decrease the overall rhythmicity with which a neuron fires, while shifting the preferred phase of spiking within an oscillatory cycle. As a result, the relatively weak input provided by tACS has the potential to powerfully alter the temporal structure of neural activity.

## Materials and Methods

### Experimental Design

To characterize the effects of tACS under conditions matching human use, we collected from two adult male rhesus monkeys (*M. mulatta*, 7 and 10 years old; 8 and 14 kg, respectively). The complete dataset includes a reanalysis of the experiments described in Vieira et al. (2020) and Krause et al. (2019) as well as new data from 157 neurons that have not been reported previously. These new data were collected using approaches and techniques that were virtually identical to our previous work, which are described in detail below. All procedures were approved by the Montreal Neurological Institute’s Animal Care Committee (Protocol #5031). The work was supervised by qualified veterinary staff, and followed the guidelines recommended by the Canadian Council on Animal Care. Animals received regular environmental enrichment, including access to a large play arena, and were socially housed when not in the laboratory. Animals were not assigned to specific experimental groups because the relevant unit of analysis are individual neurons, which were tested using a within-subject design (i.e., tACS vs baseline).

### Behavioral Task

We used a simple fixation task to ensure that animals remained in a consistent behavioral state throughout the experiment. Animals sat in a standard primate chair (Crist Instruments, Hagerstown Maryland), placed 57 cm from a computer monitor that covered the central 30° x 60° of the visual field. Eye position was monitored with an infrared eye tracker (SR Research, Ontario). The monkeys were trained to fixate a small black target (0.5°) presented against a neutral grey background (54 cd/m^2^) in exchange for juice rewards. Rewards were dispensed according to a schedule with a flat hazard function (exponential distribution with a mean of ~1 s). This arrangement minimized external factors known to influence oscillations, such as patterned visual stimulation, eye movements, and predictable rewards, allowing us to examine the effects of tACS against a consistent background. Custom software written in Matlab (The Mathworks, Natick, MA, USA) controlled the behavioral task and coordinated the eye tracker, tES stimulator, and recording hardware.

### Transcranial Alternating Current Stimulation

We applied tACS in a manner designed to closely mimic human use. To plan the stimulation, individualized finite-element models were built from preoperative MRIs of each animal’s head and neck (*66*). These were solved to identify electrode configurations that optimally stimulated neurons at our V4 and hippocampal recording sites. The hippocampal configurations produced an average field strength of 0.2 V/m (max: 0.35 V/m) in the hippocampus during ± 2 mA stimulation (*9*). Since V4 is on the cortical surface, the electric field therein was stronger: ~1 V/m at ±1 mA (*10*). These conditions therefore bracket the best available estimates for field strengths achievable with two electrode-montages in humans. (*67, 68*). Our modelling approach also ensured that the brain implants required for these experiment do not produce abnormally strong electric fields elsewhere in the brain.

The stimulation was delivered through 1-cm “high-definition” Ag/AgCl electrodes (PISTIM; Neuroelectrics; Barcelona, Spain). These were coated with a conductive gel (SignaGel; Parker Laboratories, Fairchild New Jersey) and attached to the animals’ intact scalps with a thin layer of Kwik-Sil (World Precision Instruments), a biocompatible silicon elastomer. Electrode impedance was typically between 1 and 2 kΩ, and always below 10 kΩ to prevent skin damage. In most experiments, current was applied via an unmodified StarStim8 system (Neuroelectrics, Barcelona, Spain). The stimulation waveform consisted of a 1.5- or 5-minute sinewave of the specified frequency (5 or 20 Hz) and amplitude (±1 or ±2 mA). At the beginning of each stimulation block, current was linearly ramped up from 0 to the maximum intensity over five seconds; this process was repeated in reverse at the end of each block. This minimizes the sensations produced by the onset and offset of stimulation. To measure baseline entrainment, “sham” stimulation repeated the initial and final ramps, without delivering current during the middle of the block. In a few sessions, a DC-STIMULATOR PLUS (NeuroConn; Munich, Germany), driven in remote mode by a Model 4053B signal generator (BK Precision, Yorba Linda California), was used to deliver stimulation instead. This equipment could not produce the slow ramps, so current was immediately switched on and off instead in these sessions. No significant differences between these sessions in terms of PLV change (p > 0.31; Z=1.01; Wilcoxon rank-sum test) or proportion of neurons affected (p > 0.1; chi2 = 2.83) was observed. Baseline and tACS blocks were separated by an intertrial interval of 1.5 or 5 minutes.

Although stimulation of peripheral afferents in the retina and skin can potentially confound behavioral experiments, we have previously demonstrated that neither likely account for the neural effects reported here (*9, 10*).

### Electrophysiological Recording

The target structures were accessed through sterile plastic recording chambers implanted on the skull (Crist Instruments; Hagerstown, Maryland). After penetrating the dura with a 22-gauge stainless steel guide tube, we lowered 32-channel linear arrays (V-Probe; Plexon: Dallas, Texas) into the target with a NAN Microdrive (NAN Instruments; Nazareth Illit, Israel). The location and depths of the targeted structures were confirmed via a post-operative CT (hippocampus) or MRI (V4).

The raw signals from each electrode were digitized by a Ripple Neural Interface Processor (Ripple Neuro: Salt Lake City, Utah). During acquisition, the signals were band-pass filtered between 0.3 and 7500 Hz, digitized at 0.5μV/16-bit resolution, and stored at 30,000 Hz for subsequent analysis. The artifacts from tACS stimulation did not exceed the amplifier’s linear range (±12 mV), allowing us to continuously record spiking and LFP signals during stimulation. Single units were identified offline by bandpass filtering the signal between 0.5 and 7 kHz with a third-order Butterworth filter. Spikes were initially detected as crossings exceeding a threshold of ±3 standard deviations, robustly estimated for each channel. Short segments around each threshold crossing were then extracted and clustered with UltraMegaSort 2000, a *k-*means based overclustering algorithm (*69*). Units were manually reviewed to ensure they had a consistent width and amplitude, a clear refractory period, and good separation in PCA space. Loss of signal during parts of the tACS cycle could produce spurious changes in entrainment, but control analyses for the same equipment and analysis pipeline suggest this is unlikely (e.g., Figure S2 of *9*).

### Phase-locking Analysis

We quantified neural entrainment by calculating pairwise phase consistency (PPC) values for each cell, a measure of the synchronization between the phase of an on-going signal (here, the LFP or tACS) and a point process (spiking activity). Although computationally intensive, this method has several statistical advantages over other common measures of phase-locking (*70*).

These were calculated using spikes obtained from one channel and the continuous signal from an adjacent channel (150 μm away), so as to avoid spectral contamination that may artifactually inflate measures of entrainment (*71*). Using this local signal, rather than measuring entrainment to copy of the tACS output, also ensures that referencing remained constant across conditions and accounts for any physiological distortion of the tACS waveform (*72*). To define the phase, the wideband signal was filtered into a ±1 Hz range around the frequency of interest (i.e., 4—6 or 19—21 Hz). The instantaneous phase at the time of each spike was derived from its Hilbert transform and used to calculate the PPC. PPCs were calculated for each condition separately and compared across conditions via a randomization test. For presentation purposes, these were then converted to phase-locking values (PLVs), a more commonly used measure of spike entrainment. Conveniently, PLVs are also equivalent (under some simplifying assumptions) to spike-field coherence, another oft-used metric of spike-LFP coupling.

In the Results section, we focus only on the frequency bands of interest (5±1 or 20±1 Hz) because the effects of tACS were confined to a narrow window around the stimulation frequencies. PLV values calculated in ±1 Hz bands from 2-100 Hz revealed significantly increased entrainment only in the 2-7 Hz range. Likewise, cells that showed decreased entrainment did not, on average, become coupled to any other frequency component during stimulation (all p > 0.05; Wilcoxon sign-rank tests).

Our estimates of oscillation phase were relative to the electric potential in the extracellular space near each neuron. For the baseline condition, this reflects the LFP, while for the tACS condition it is dominated by the instantaneous phase of tACS as detected by our recording electrode. Thus, the observed differences in the preferred phase of spiking cannot be attributed to a difference in referencing conditions (*73*), as these were identical for baseline and tACS conditions. Moreover, the change in phase observed with tACS cannot be attributed simply to phase differences between tACS and ongoing oscillations. Given the open-loop nature of our experiments, this difference was random from block to block, but we observed a highly consistent preference change between baseline and tACS blocks.

### Cell Type Identification

Labelling individual neurons from which data were collected is technically infeasible in nonhuman primates, so putative interneurons were identified based on the shape of their extracellular action potential. Many interneurons contain Kv3 potassium channels, whose fast deactivation rates help quickly repolarize the cell, causing the action potentials to become thinner than those produced by more sluggish channels present in excitatory cells (*74*). The trough-to-peak width of the extracellular action potential has therefore been proposed as a method for separating putative inhibitory and excitatory cells (*75*). For each neuron, we measured the trough-to-peak width on its average waveform, using spikes collected in all conditions. Our data exhibited a clearly bimodal distribution (Figure 6A), and we applied a hard threshold at 250 μs (*76*), classifying neurons with narrower spikes as putative interneurons and those with broader spikes as putative excitatory cells. Unsupervised clustering with a Gaussian mixture model yielded very similar classifications.

A more fine-grained categorization of cell types may be possible if additional features of the extracellular action potential are included in the clustering analysis (*77*). However, it remains unclear how these functional categories map onto specific morphologies. On the other hand, the two-way classification used here allows us to test a specific hypothesis about the relationship between cell shape and determine if interneurons are less affected by exogenous electric fields.

However, not every narrow-spiking neuron in our data set is likely to be a fast-spiking GABAeric interneuron. Previous work has identified neurons that emit narrow action potentials despite having spiny, pyramidal morphology (*78*) and, in primates, up to 25% of Kv3-expressing neurons are non-GABAergic (*79*). An alternate explanation for our findings in Figure 6 is that these non-GABAergic narrow-spiking neurons are affected by tACS, while the “true” interneurons are unresponsive. However, the proportion of affected cells in our data (23/51) is significantly higher than would be expected under a 25%/75% split (p < 0.01; χ(1) = 10.99), suggesting that this is unlikely to occur in our data.

### Model Simulations

The Stuart-Landau oscillator described above consists of two ordinary differential equations with 3 parameters (*36*). The frequency of the ongoing oscillation is determined by the ω parameter, while its stability is controlled by *λ:* values above zero produce self-sustaining oscillations, while setting *λ* < 0 causes the oscillations to decay with time. Finally, *γ* dampens the system, controlling how quickly it settles into a steady-state amplitude.

The model equations were solved numerically using Matlab’s `ode45` function, an explicit Runge-Kutta (4,5) method, using the solver’s default parameters and the initial conditions *x* = 0, *y* = −1. The model was run for 200 s total. In stimulation conditions, the external drive *s(t)* was included only after 40 s so that the model could reach steady-state. In both cases, we defined the amplitude as √*2* times root-mean-squared amplitude of the signal; this corresponds to an amplitude of 1 for a standard sine wave but corrects for any transients created by numerical integration. To generate Figure 4C, we evaluated all combinations of phase mismatch (0-315°, in steps of 45°) and frequency mismatch (50-150% of the baseline oscillation’s cycle, in steps of 10%), for values of k ranging from 0 to 100% of the baseline oscillation’s amplitude. These were then numerically integrated with Matlab’s `trapz` to find the net effect across phase and frequency shown in Figure 4D. Similar results were obtained using Julia and its `Tsit5` solver.

As the model does not have explicit physical units, we performed a limited exploration of the parameter space, to determine whether a qualitative match to our data could be found. For Figure 4, the values were Λ=0.2, ω=0.5, and Λ=1.0, which were similar to those used in a previous study of the model (*36*). However, these precise parameters were not required to reproduce the negative flanks shown in Figure 4C and 4D. We ran 250 additional simulations, setting Λ, ω, and *λ* independently to random values between 0.1 and 10; smaller values took prohibitively long to reach steady-state. In more than half of these simulations (162/250), at least one value of k (10-150%, in steps of 10%) reduced the total entrainment, as in Figure 4D. Indeed, the “quenching” behavior we have observed is actually a universal feature of many kinds of oscillators (*80*), and as shown in the Supplemental Analysis, similar results can be derived analytically from a simplified version of this model (*81*).

### Quantification and Statistical Analysis

Where possible, statistical analyses were performed using nonparametric tests that avoid distributional assumptions; population-level analyses were carried using Wilcoxon rank-sum and sign-rank tests, as appropriate. We used 95 percent confidence intervals of the median as population dispersion, calculated using the formula in (*82*). All statistical tests are two-tailed, using sample sizes derived from our previous work. Significance of differences between categories was assessed using the chi-square test. Sample sizes were determined based on our previous work and data were analyzed using Matlab (The Mathworks, Natick, MA, USA), the CircStats toolbox (*83*), and Julia (*84*). As we reviewed in (*9*), the effect sizes reported here are similar to those produced by changes in behavioral state and sensory input, and therefore are likely to be physiologically relevant.

## Data and Code Availability

The data that support the findings of this study are available from the corresponding author upon reasonable request.

## Acknowledgments

We thank Julie Coursol, Cathy Hunt, and Dr. Fernando Chaurand for outstanding technical assistance. Melanie Segado provided helpful comments on the paper. This work was supported by the Canadian Institutes of Health Research Grant MOP-115178 (CCP), Parkinson’s Canada Pilot Project Grant PPG-2020-0000000033 (MRK), and Natural Sciences and Engineering Council of Canada Discovery Grant #210977 (JPT)

## Author Contributions

Conceptualization: MRK, PGV, CCP

Methodology: MRK, PGV, CCP

Investigation: MRK, PGV

Formal Analysis: MRK, PGV, JPT, CCP

Visualization: MRK, PGV

Supervision: CCP

Writing—original draft: MRK, PGV, JPT, CCP

Writing—review & editing: MRK, PGV, JPT, CCP

## Supplementary Materials

### Supplementary Text

#### Asynchronous Quenching in Simplified Stuart-Landau Oscillators

We start with a system of two coupled Stuart-Landau oscillators

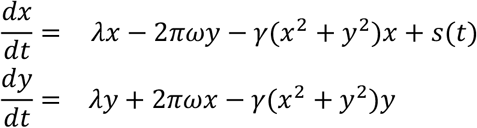

where s(t) is a periodic forcing of the form s(t) = k · sin(2πω_s_ t + ϕ_s_), where k is its amplitude and ω_s_ is the frequency of the stimulation. The central result of the model presented in the paper (Figure 4) is that of “asynchronous quenching”: entrainment is maximal when the frequency of the forcing ω_s_ is closest to the intrinsic frequency of the oscillators ω. This result cannot be explained by a simple summation of periodic oscillations corresponding to the intrinsic and driven waves, regardless of their phase offset. To gain a mechanistic understanding of this finding, it is worthwhile to consider a simplified linear system obtained by setting γ = 0, which reduces the model to

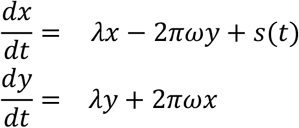

When k = 0, the eigenvalues of the above expressions are λ ± iω, so setting λ < 0 ensures that the system is globally stable. Its dynamics are therefore easy to track analytically, but the behavior of the model is simplified: there are no reductions in entrainment as in Figure 4C and 4D. We can derive the maximal amplitude of the forced oscillator in this simplified expression (Mori and Kuramoto, 1998):

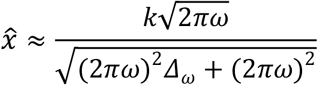

where Δ_ω_ = (2πω – 2πω_s_)^2^. Here, the numerator reflects the amplitude and frequency of the forced oscillation, while the denominator takes into account the difference between the intrinsic and forced oscillation’s frequencies. As an example, consider numerical simulation where a 2Hz forced oscillation is applied that matched the intrinsic frequency (i.e., ω = ω_s_ = 2 Hz and therefore Δ_w_ = 0). In this case, the response of *x(t)* is prominent (Figure S4, top row). However, with unmatched frequencies (ω_s_ = 1 Hz or ω_s_ = 3 Hz), the response is attenuated. Interestingly, similar results are obtained even when the intrinsic oscillation is left to vanish before the forced oscillation is applied (Figure S4, bottom row). With λ = —1, Figure S4A shows agreement between the numerical and analytical results. This is robust against changes in phase, which alters the shape of the response without compromising the fit obtained from 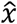, as shown in Figure S5B. Finally, a similar distribution is obtained with λ = 0.1, though in this case the system’s eigenvalues are positive and therefore the amplitude of *x(t)* is markedly higher (Figure S5C) and not amenable to the approximation described above. In all cases shown in Figure S5, the range of forced frequencies leading to a change in the mean amplitude of *x* depends on *k*, with larger values broadening this range. Finally, asynchronous quenching can also be obtained in coupled oscillators where a time delay is introduced, as shown in Figure S6.

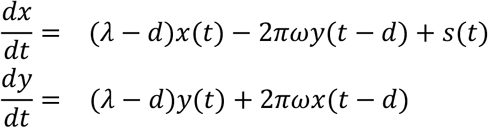

**Fig. S1.**
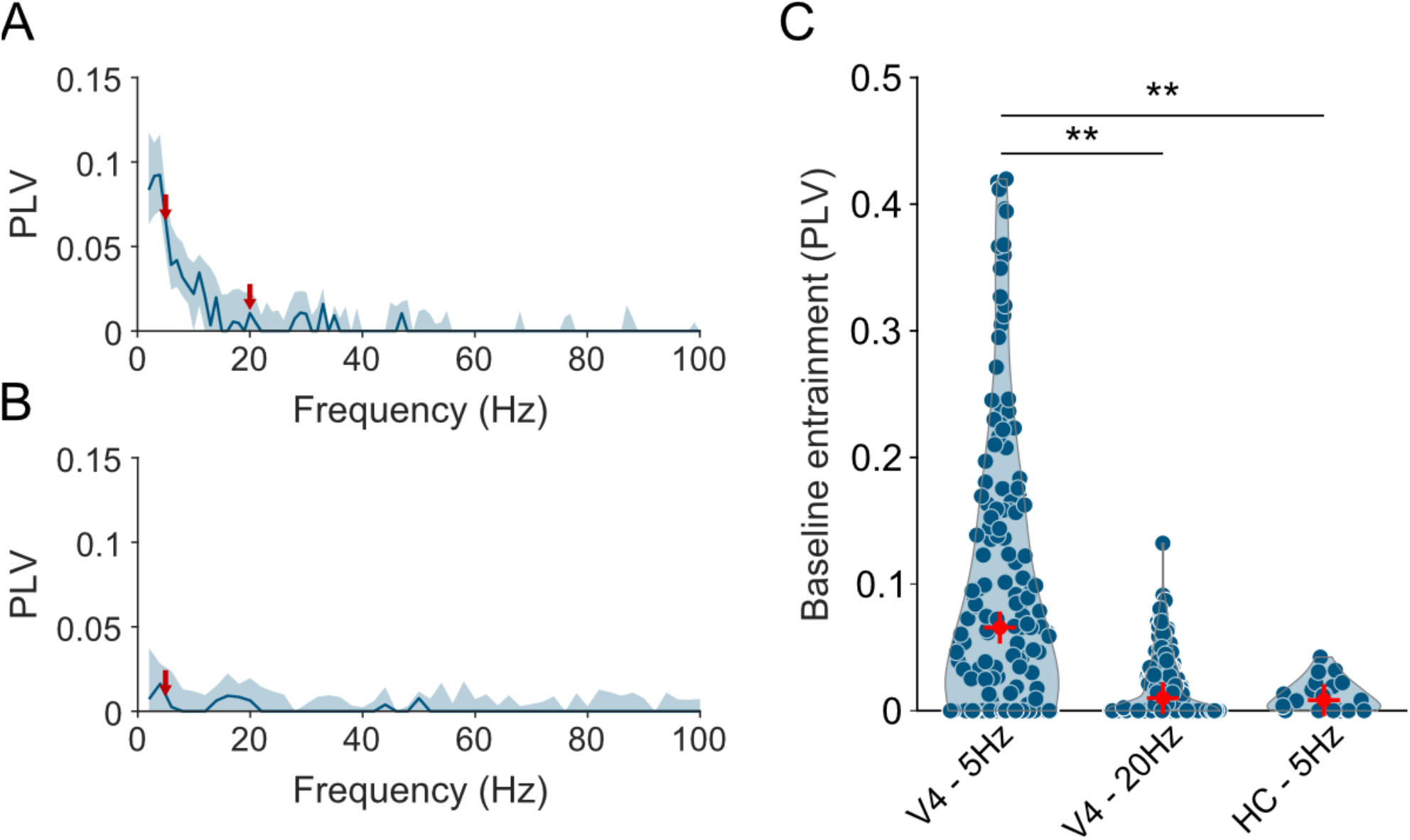
Baseline entrainment of neurons in each experimental condition. **(A)** Entrainment spectra for V4 neurons (N=157), showing phase-locking to the local field potential in ± 1 Hz frequency bands. The median and 95% confidence interval are shown, calculated from all V4 neurons (i.e., those used in subsequent 5 Hz and 20 Hz tACS experiments). Red arrows indicate the tACS frequencies used in those experiments. **(B)** Entrainment spectra for hippocampal neurons (N=21), plotted in the same style. **(C)** Individual baseline values for each cell in the three experiment conditions. The red cross indicates the median and ** indicates significance at the p < 0.01 level. See also Main Figure 1; note that the V4 data is divided across Panels A and B of Main Figure 1.

**Fig. S2.**
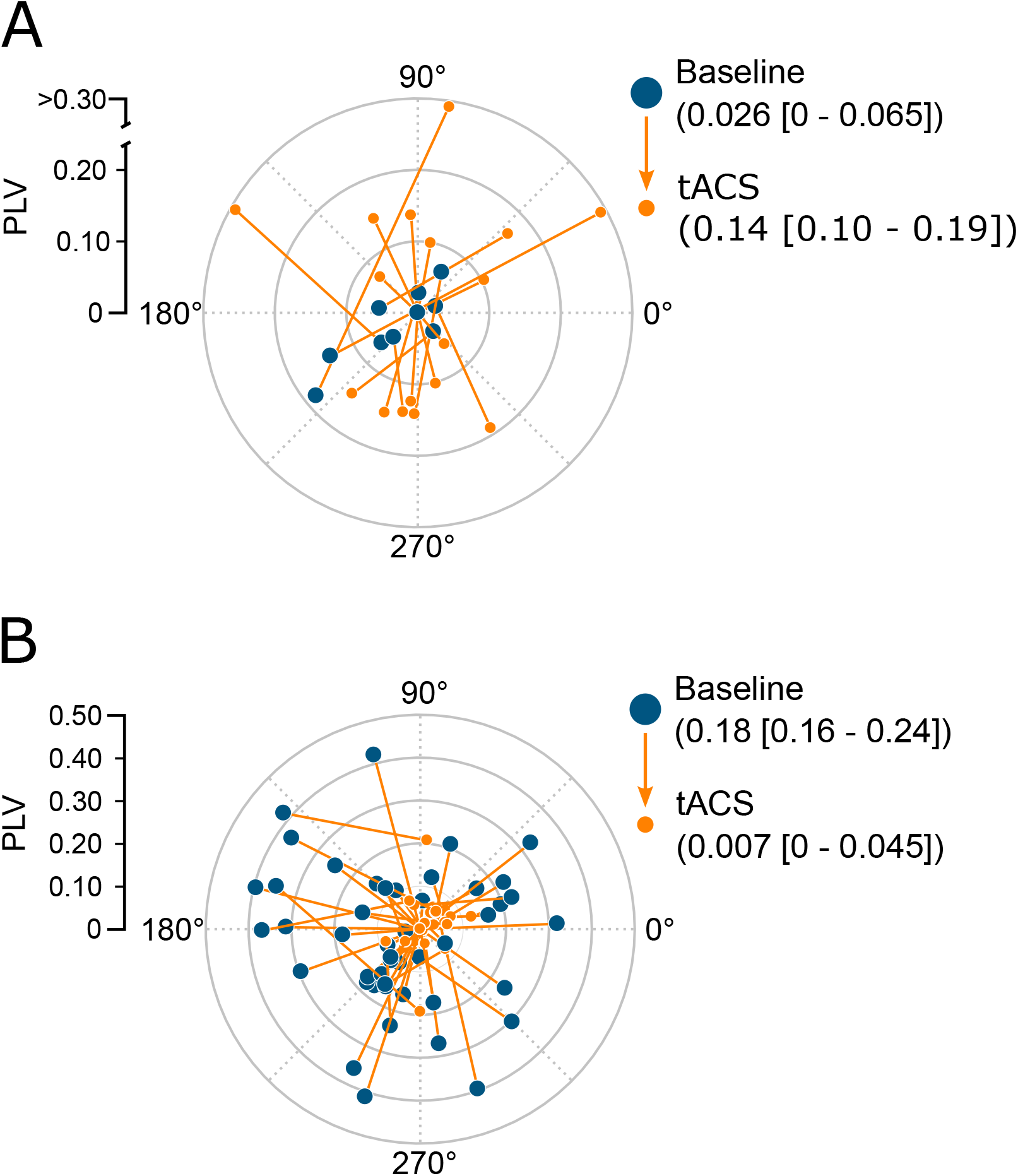
Polar vector fields for neurons whose entrainment was altered by tACS. Polar plots summarizing the combined effects of tACS on entrainment strength (PLV, eccentric direction) and entrainment phase (polar angle) on neurons that became significantly more (A) or less (B) entrained by tACS. See also Main Figure 2.

**Fig. S3.**
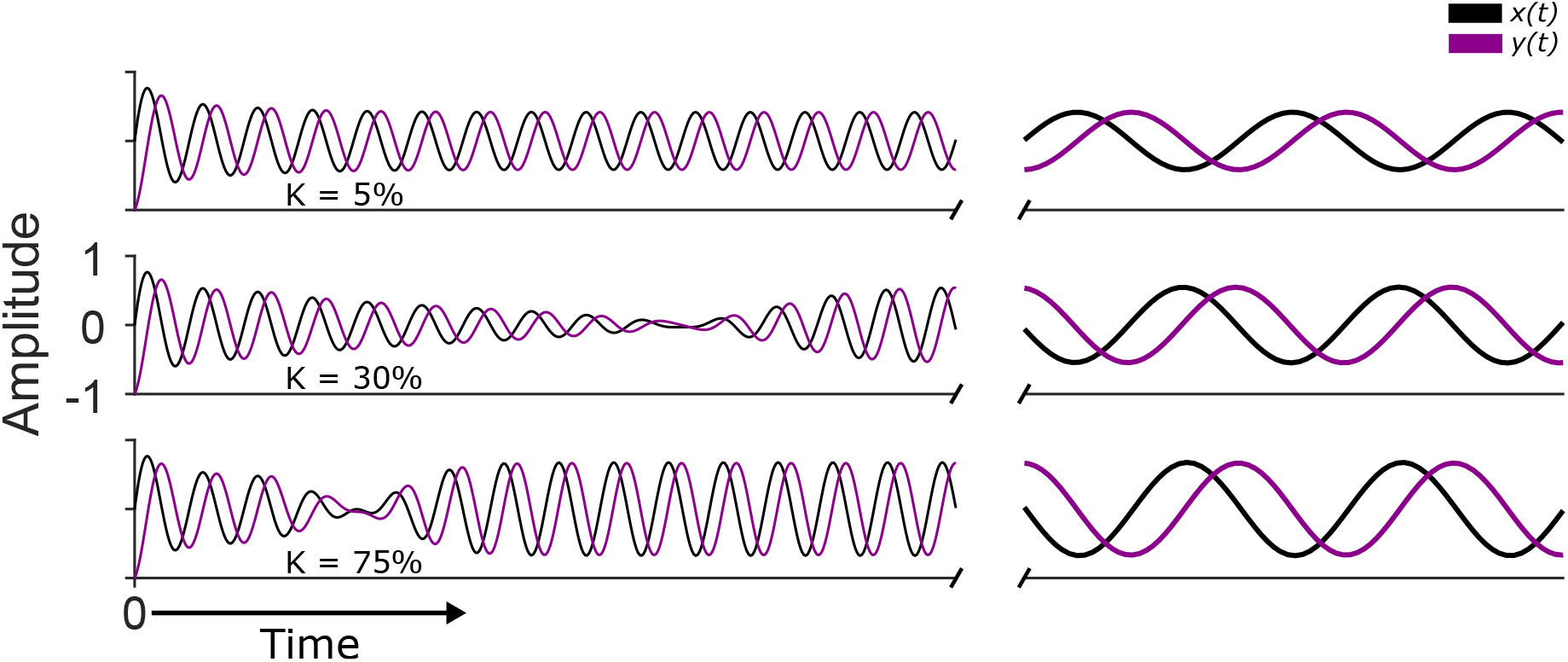
Both model populations are affected similarly by tACS. The values of *x* (black) and *y* (magenta) during three runs of the model with *k* = 5, 30, and 75 percent, demonstrating that they are phase-shifted copies of each other. Note that baseline is not shown here. See also Main Figure 4.

**Fig. S4.**
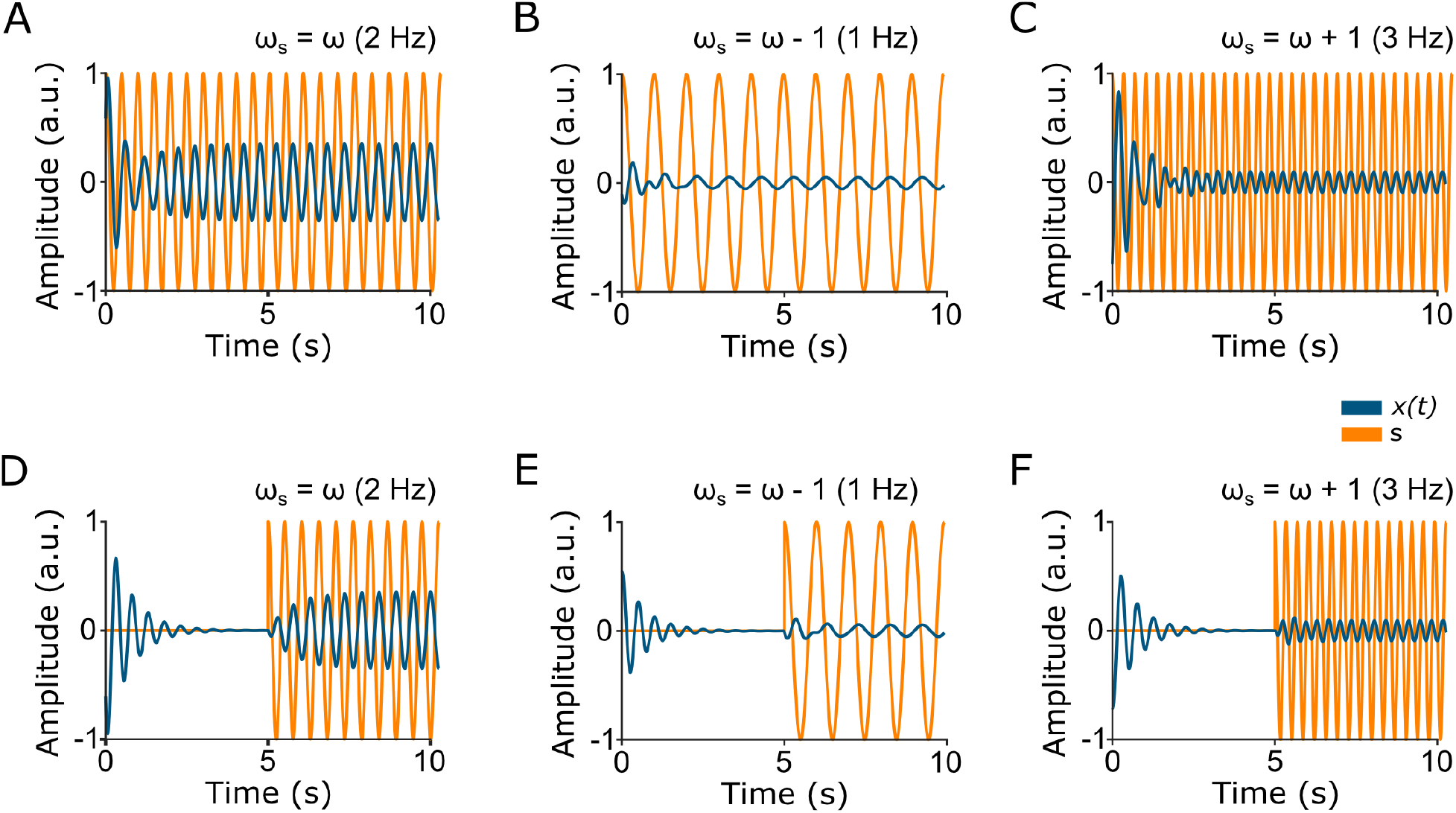
Driving a Stuart-Landau system with a forced oscillation. In all panels, *ω* was set to 2 Hz while the frequency of stimulation *ω_s_* was varied. **(A-C)** Stimulation (orange line) was applied to an ongoing oscillation (blue) at the same frequency (A), at a slightly lower frequency (B), and at a slightly higher frequency (C). **(D-F)** As above, except that the ongoing oscillation was allowed to decay before stimulation was applied.

**Fig. S5.**
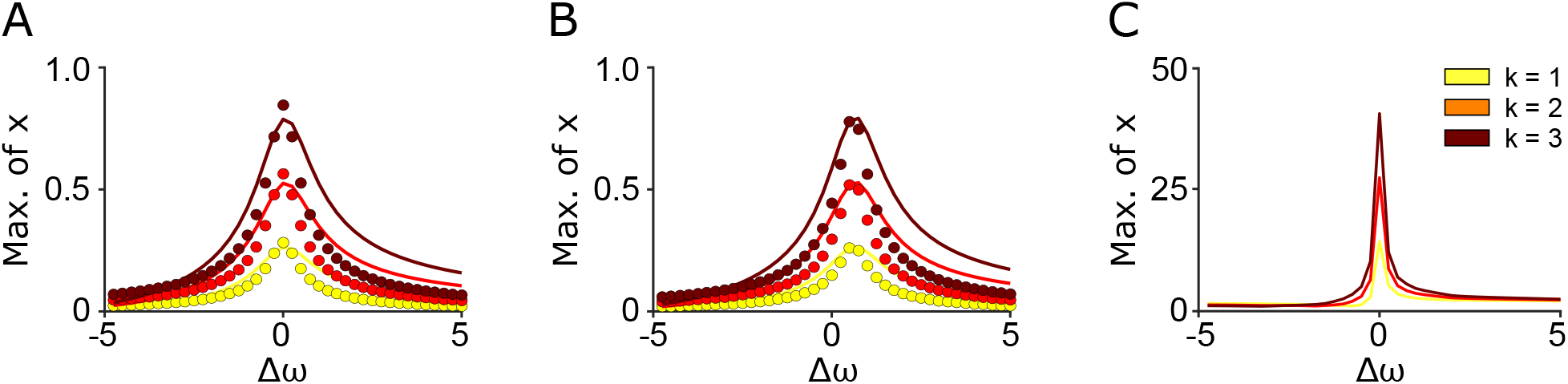
Comparison of numerical and analytical results. **(A-B)** Filled circles indicate the analytical solution 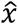, while the lines are derived from the numerical simulation. In all panels the intrinsic frequency was set to 5 Hz with a phase shift of the forced oscillation set to zero (A) or 1.5 radians (B). **(C)** Similar results were obtained from an unstable model where *λ* = 0.1.

**Fig. S6.**
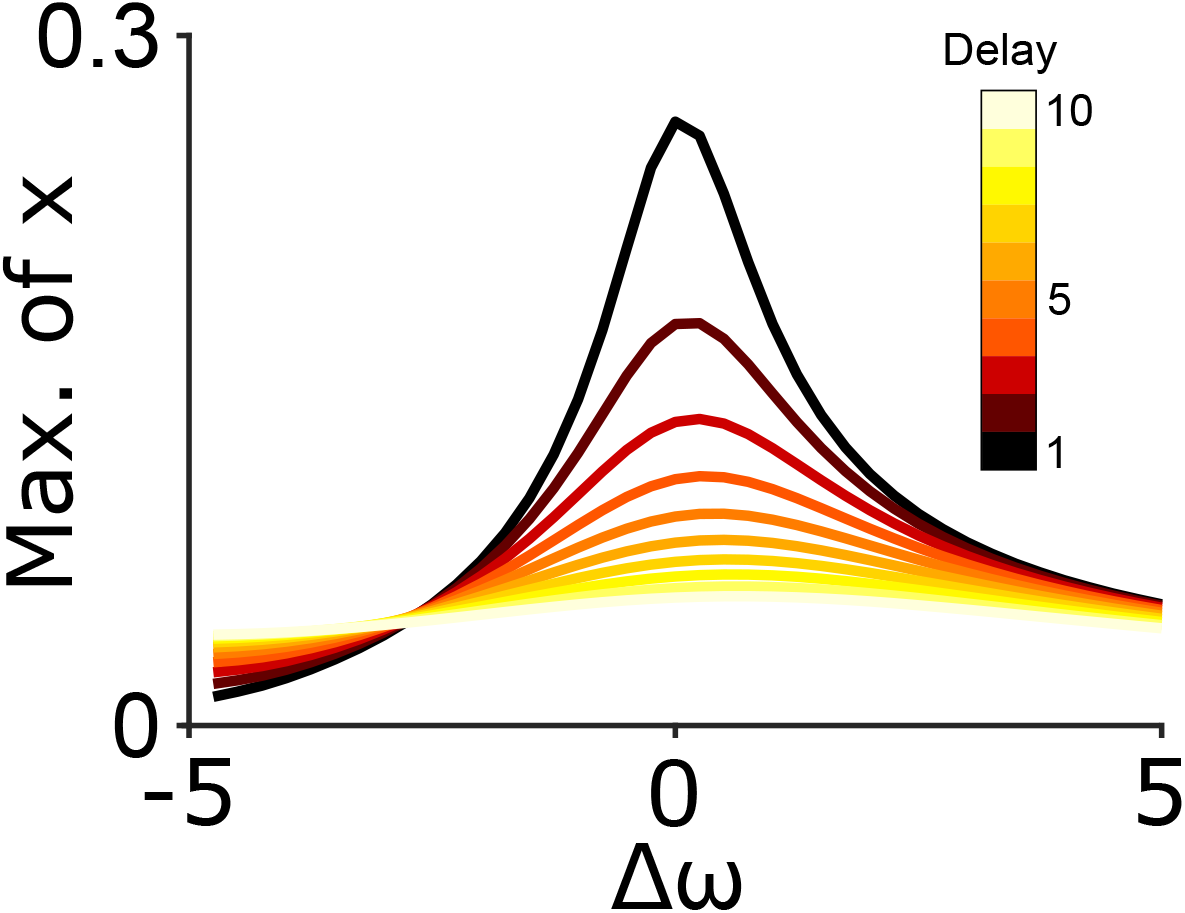
Asynchronous quenching is robust to delays between the two oscillators. Results of simulations where the delays ranged from 1 (black) to 10 (yellow).

## References

1. P. Fries, Rhythms for Cognition: Communication through Coherence. Neuron 88, 220–235 (2015).

2. D. J. Mogul, W. van Drongelen, Electrical control of epilepsy. Annu Rev Biomed Eng 16, 483–504 (2014).

3. S. Grover, J. A. Nguyen, R. M. G. Reinhart, Synchronizing Brain Rhythms to Improve Cognition. Annu Rev Med 72, 29–43 (2021).

4. C. Y. Chan, C. Nicholson, Modulation by applied electric fields of Purkinje and stellate cell activity in the isolated turtle cerebellum. J Physiol 371, 89–114 (1986).

5. C. A. Anastassiou, R. Perin, H. Markram, C. Koch, Ephaptic coupling of cortical neurons. Nat Neurosci 14, 217–223 (2011).

6. F. Frohlich, D. A. McCormick, Endogenous electric fields may guide neocortical network activity. Neuron 67, 129–143 (2010).

7. S. Ozen et al., Transcranial electric stimulation entrains cortical neuronal populations in rats. J Neurosci 30, 11476–11485 (2010).

8. L. Johnson et al., Dose-dependent effects of transcranial alternating current stimulation on spike timing in awake nonhuman primates. Sci Adv 6, (2020).

9. M. R. Krause, P. G. Vieira, B. A. Csorba, P. K. Pilly, C. C. Pack, Transcranial alternating current stimulation entrains single-neuron activity in the primate brain. Proc Natl Acad Sci U S A 116, 5747–5755 (2019).

10. P. G. Vieira, M. R. Krause, C. C. Pack, tACS entrains neural activity while somatosensory input is blocked. PLoS Biol 18, e3000834 (2020).

11. A. Coldea, S. Morand, D. Veniero, M. Harvey, G. Thut, Parietal alpha tACS shows inconsistent effects on visuospatial attention. PLoS One 16, e0255424 (2021).

12. F. H. Kasten, K. Duecker, M. C. Maack, A. Meiser, C. S. Herrmann, Integrating electric field modeling and neuroimaging to explain inter-individual variability of tACS effects. Nat Commun 10, 5427 (2019).

13. D. Veniero, C. S. Y. Benwell, M. M. Ahrens, G. Thut, Inconsistent Effects of Parietal alpha-tACS on Pseudoneglect across Two Experiments: A Failed Internal Replication. Front Psychol 8, 952 (2017).

14. F. H. Kasten, C. S. Herrmann, The hidden state-dependency of transcranial alternating current stimulation (tACS). bioRxiv, 2020.2012.2023.423984 (2020).

15. S. Alagapan et al., Modulation of Cortical Oscillations by Low-Frequency Direct Cortical Stimulation Is State-Dependent. PLoS Biol 14, e1002424 (2016).

16. M. Feurra et al., State-dependent effects of transcranial oscillatory currents on the motor system: what you think matters. J Neurosci 33, 17483–17489 (2013).

17. T. Neuling, S. Rach, C. S. Herrmann, Orchestrating neuronal networks: sustained aftereffects of transcranial alternating current stimulation depend upon brain states. Front Hum Neurosci 7, 161 (2013).

18. P. Ruhnau et al., Eyes wide shut: Transcranial alternating current stimulation drives alpha rhythm in a state dependent manner. Sci Rep 6, 27138 (2016).

19. A. K. Chiang, C. J. Rennie, P. A. Robinson, S. J. van Albada, C. C. Kerr, Age trends and sex differences of alpha rhythms including split alpha peaks. Clin Neurophysiol 122, 1505–1517 (2011).

20. K. J. Mathewson et al., Regional EEG alpha power, coherence, and behavioral symptomatology in autism spectrum disorder. Clin Neurophysiol 123, 1798–1809 (2012).

21. S. Okuhata, T. Kusanagi, T. Kobayashi, Parietal EEG alpha suppression time of memory retrieval reflects memory load while the alpha power of memory maintenance is a composite of the visual process according to simultaneous and successive Sternberg memory tasks. Neurosci Lett 555, 79–84 (2013).

22. O. Jensen, J. Gelfand, J. Kounios, J. E. Lisman, Oscillations in the alpha band (9-12 Hz) increase with memory load during retention in a short-term memory task. Cereb Cortex 12, 877–882 (2002).

23. C. S. Herrmann, S. Rach, T. Neuling, D. Struber, Transcranial alternating current stimulation: a review of the underlying mechanisms and modulation of cognitive processes. Front Hum Neurosci 7, 279 (2013).

24. D. Reato, A. Rahman, M. Bikson, L. C. Parra, Low-intensity electrical stimulation affects network dynamics by modulating population rate and spike timing. J Neurosci 30, 15067–15079 (2010).

25. J. K. Deans, A. D. Powell, J. G. Jefferys, Sensitivity of coherent oscillations in rat hippocampus to AC electric fields. J Physiol 583, 555–565 (2007).

26. M. Voroslakos et al., Direct effects of transcranial electric stimulation on brain circuits in rats and humans. Nat Commun 9, 483 (2018).

27. A. Berenyi, M. Belluscio, D. Mao, G. Buzsaki, Closed-loop control of epilepsy by transcranial electrical stimulation. Science 337, 735–737 (2012).

28. S. L. Schmidt, A. K. Iyengar, A. A. Foulser, M. R. Boyle, F. Frohlich, Endogenous cortical oscillations constrain neuromodulation by weak electric fields. Brain Stimul 7, 878–889 (2014).

29. G. Spyropoulos, C. A. Bosman, P. Fries, A theta rhythm in macaque visual cortex and its attentional modulation. Proc Natl Acad Sci U S A 115, E5614–E5623 (2018).

30. P. D. Oldham, A note on the analysis of repeated measurements of the same subjects. J Chronic Dis 15, 969–977 (1962).

31. K. Kar, J. Duijnhouwer, B. Krekelberg, Transcranial Alternating Current Stimulation Attenuates Neuronal Adaptation. J Neurosci 37, 2325–2335 (2017).

32. B. Voloh, M. Oemisch, T. Womelsdorf, Phase of firing coding of learning variables across the fronto-striatal network during feature-based learning. Nat Commun 11, 4669 (2020).

33. E. H. Park, E. Barreto, B. J. Gluckman, S. J. Schiff, P. So, A model of the effects of applied electric fields on neuronal synchronization. J Comput Neurosci 19, 53–70 (2005).

34. T. Radman, R. L. Ramos, J. C. Brumberg, M. Bikson, Role of cortical cell type and morphology in subthreshold and suprathreshold uniform electric field stimulation in vitro. Brain Stimul 2, 215–228, 228 e211-213 (2009).

35. D. Reato, A. Rahman, M. Bikson, L. C. Parra, Effects of weak transcranial alternating current stimulation on brain activity-a review of known mechanisms from animal studies. Front Hum Neurosci 7, 687 (2013).

36. K. B. Doelling, M. F. Assaneo, Neural oscillations are a start toward understanding brain activity rather than the end. PLoS Biol 19, e3001234 (2021).

37. W. R. Softky, C. Koch, The highly irregular firing of cortical cells is inconsistent with temporal integration of random EPSPs. J Neurosci 13, 334–350 (1993).

38. F. Mansouri, K. Dunlop, P. Giacobbe, J. Downar, J. Zariffa, A Fast EEG Forecasting Algorithm for Phase-Locked Transcranial Electrical Stimulation of the Human Brain. Front Neurosci 11, 401 (2017).

39. S. P. Burns, D. Xing, R. M. Shapley, Is gamma-band activity in the local field potential of V1 cortex a “clock” or filtered noise? J Neurosci 31, 9658–9664 (2011).

40. S. R. Cole, B. Voytek, Brain Oscillations and the Importance of Waveform Shape. Trends Cogn Sci 21, 137–149 (2017).

41. A. T. Schaefer, K. Angelo, H. Spors, T. W. Margrie, Neuronal oscillations enhance stimulus discrimination by ensuring action potential precision. PLoS Biol 4, e163 (2006).

42. D. K. Bilkey, U. Heinemann, Intrinsic theta-frequency membrane potential oscillations in layer III/V perirhinal cortex neurons of the rat. Hippocampus 9, 510–518 (1999).

43. A. S. Aberra, A. V. Peterchev, W. M. Grill, Biophysically realistic neuron models for simulation of cortical stimulation. J Neural Eng 15, 066023 (2018).

44. W. A. Huang et al., Transcranial alternating current stimulation entrains alpha oscillations by preferential phase synchronization of fast-spiking cortical neurons to stimulation waveform. Nat Commun 12, 3151 (2021).

45. N. C. Venables, E. M. Bernat, S. R. Sponheim, Genetic and disorder-specific aspects of resting state EEG abnormalities in schizophrenia. Schizophr Bull 35, 826–839 (2009).

46. S. Santaniello et al., Therapeutic mechanisms of high-frequency stimulation in Parkinson’s disease and neural restoration via loop-based reinforcement. Proc Natl Acad Sci U S A 112, E586–595 (2015).

47. D. Premraj, K. Manoj, S. A. Pawar, R. I. Sujith, Effect of amplitude and frequency of limit cycle oscillators on their coupled and forced dynamics. Nonlinear Dynamics 103, 1439–1452 (2021).

48. S. Sugiyama et al., Suppression of Low-Frequency Gamma Oscillations by Activation of 40-Hz Oscillation. Cerebral Cortex, (2021).

49. Y. Hashimoto, Y. Yotsumoto, The Amount of Time Dilation for Visual Flickers Corresponds to the Amount of Neural Entrainments Measured by EEG. Front Comput Neurosci 12, 30 (2018).

50. S. Marchesotti et al., Selective enhancement of low-gamma activity by tACS improves phonemic processing and reading accuracy in dyslexia. PLoS Biol 18, e3000833 (2020).

51. Y. Cabral-Calderin, M. Wilke, Probing the Link Between Perception and Oscillations: Lessons from Transcranial Alternating Current Stimulation. Neuroscientist 26, 57–73 (2020).

52. L. D. Liu, R. M. Haefner, C. C. Pack, A neural basis for the spatial suppression of visual motion perception. Elife 5, (2016).

53. E. Zohary, S. Celebrini, K. H. Britten, W. T. Newsome, Neuronal plasticity that underlies improvement in perceptual performance. Science 263, 1289–1292 (1994).

54. M. R. Cohen, J. H. Maunsell, Attention improves performance primarily by reducing interneuronal correlations. Nat Neurosci 12, 1594–1600 (2009).

55. P. Fries, J. H. Reynolds, A. E. Rorie, R. Desimone, Modulation of oscillatory neuronal synchronization by selective visual attention. Science 291, 1560–1563 (2001).

56. B. N. Pasley, E. A. Allen, R. D. Freeman, State-dependent variability of neuronal responses to transcranial magnetic stimulation of the visual cortex. Neuron 62, 291–303 (2009).

57. E. A. Allen, B. N. Pasley, T. Duong, R. D. Freeman, Transcranial magnetic stimulation elicits coupled neural and hemodynamic consequences. Science 317, 1918–1921 (2007).

58. M. L. Waterston, C. C. Pack, Improved discrimination of visual stimuli following repetitive transcranial magnetic stimulation. PLoS One 5, e10354 (2010).

59. W. Klimesch, alpha-band oscillations, attention, and controlled access to stored information. Trends Cogn Sci 16, 606–617 (2012).

60. S. E. Ehrhardt, H. L. Filmer, Y. Wards, J. B. Mattingley, P. E. Dux, The influence of tDCS intensity on decision-making training and transfer outcomes. J Neurophysiol 125, 385–397 (2021).

61. V. Moliadze, D. Atalay, A. Antal, W. Paulus, Close to threshold transcranial electrical stimulation preferentially activates inhibitory networks before switching to excitation with higher intensities. Brain Stimul 5, 505–511 (2012).

62. M. Okun et al., Diverse coupling of neurons to populations in sensory cortex. Nature 521, 511–515 (2015).

63. M. Witkowski et al., Mapping entrained brain oscillations during transcranial alternating current stimulation (tACS). Neuroimage 140, 89–98 (2016).

64. R. F. Helfrich et al., Entrainment of brain oscillations by transcranial alternating current stimulation. Curr Biol 24, 333–339 (2014).

65. N. K. Logothetis, The underpinnings of the BOLD functional magnetic resonance imaging signal. The Journal of neuroscience: the official journal of the Society for Neuroscience 23, 3963–3971 (2003).

66. A. Datta, Krause, M.R., Pilly, P.K., Choe, J., Zanos, T.P., Thomas, C., and Pack, C.C., On comparing in vivo intracranial recordings in non-human primates to predictions of optimized transcranial electrical stimulation. Proceedings of the 38th Annual International Conference of the IEEE Engineering in Medicine and Biology Society, (2016).

67. M. P. Jackson et al., Animal models of transcranial direct current stimulation: Methods and mechanisms. Clin Neurophysiol 127, 3425–3454 (2016).

68. Y. Huang et al., Measurements and models of electric fields in the in vivo human brain during transcranial electric stimulation. Elife 6, (2017).

69. D. N. Hill, S. B. Mehta, D. Kleinfeld, Quality metrics to accompany spike sorting of extracellular signals. J Neurosci 31, 8699–8705 (2011).

70. M. Vinck, M. van Wingerden, T. Womelsdorf, P. Fries, C. M. Pennartz, The pairwise phase consistency: a bias-free measure of rhythmic neuronal synchronization. Neuroimage 51, 112–122 (2010).

71. T. P. Zanos, P. J. Mineault, C. C. Pack, Removal of spurious correlations between spikes and local field potentials. J Neurophysiol 105, 474–486 (2011).

72. N. Noury, J. F. Hipp, M. Siegel, Physiological processes non-linearly affect electrophysiological recordings during transcranial electric stimulation. Neuroimage 140, 99–109 (2016).

73. V. Shirhatti, A. Borthakur, S. Ray, Effect of Reference Scheme on Power and Phase of the Local Field Potential. Neural Comput 28, 882–913 (2016).

74. B. Rudy et al., Contributions of Kv3 channels to neuronal excitability. Ann N Y Acad Sci 868, 304–343 (1999).

75. J. F. Mitchell, K. A. Sundberg, J. H. Reynolds, Differential attention-dependent response modulation across cell classes in macaque visual area V4. Neuron 55, 131–141 (2007).

76. M. Vinck, T. Womelsdorf, E. A. Buffalo, R. Desimone, P. Fries, Attentional modulation of cell-class-specific gamma-band synchronization in awake monkey area v4. Neuron 80, 1077–1089 (2013).

77. C. Trainito, C. von Nicolai, E. K. Miller, M. Siegel, Extracellular Spike Waveform Dissociates Four Functionally Distinct Cell Classes in Primate Cortex. Curr Biol 29, 2973–2982 e2975 (2019).

78. L. G. Nowak, R. Azouz, M. V. Sanchez-Vives, C. M. Gray, D. A. McCormick, Electrophysiological classes of cat primary visual cortical neurons in vivo as revealed by quantitative analyses. J Neurophysiol 89, 1541–1566 (2003).

79. C. M. Constantinople, A. A. Disney, J. Maffie, B. Rudy, M. J. Hawken, Quantitative analysis of neurons with Kv3 potassium channel subunits, Kv3.1b and Kv3.2, in macaque primary visual cortex. J Comp Neurol 516, 291–311 (2009).

80. A. Pikovsky, J. Kurths, M. Rosenblum, J. Kurths, Synchronization: a universal concept in nonlinear sciences. (Cambridge university press, 2003).

81. H. Mori, Y. Kuramoto, Dissipative Structures and Chaos. (Iwanami Shoten, Publishers, Tokyo, 1998).

82. M. J. Campbell, M. J. Gardner, Calculating confidence intervals for some non-parametric analyses. Br Med J (Clin Res Ed) 296, 1454–1456 (1988).

83. P. Berens, CircStat: A MATLAB Toolbox for Circular Statistics. Journal of Statistical Software 31, 1–21 (2009).

84. C. Rackauckas, Q. Nie, DifferentialEquations.jl – A Performant and Feature-Rich Ecosystem for Solving Differential Equations in Julia. Journal of Open Research Software 5, (2017).

